# Unique and Common Agonists Activate the Insect Juvenile Hormone Receptor and the Human AHR

**DOI:** 10.1101/2024.01.03.574093

**Authors:** David Sedlak, Roman Tuma, Jayaprakash Narayana Kolla, Raveendra Babu Mokhamatam, Liliia Bahrova, Michaela Lisova, Lenka Bittova, Marek Jindra

## Abstract

Transcription factors of the bHLH-PAS family play vital roles in animal development, physiology, and disease. Two members of the family require binding of low-molecular weight ligands for their activity: the vertebrate aryl hydrocarbon receptor (AHR) and the insect juvenile hormone receptor (JHR). In the fly *Drosophila melanogaster*, the paralogous proteins GCE and MET constitute the ligand-binding component of JHR complexes. Whilst GCE/MET and AHR are phylogenetically heterologous, their mode of action is similar. JHR is targeted by several synthetic agonists that serve as insecticides disrupting the insect endocrine system. AHR is an important regulator of human endocrine homeostasis and it responds to environmental pollutants and endocrine disruptors. Whether AHR signaling is affected by compounds that can activate JHR has not been reported. To address this question, we screened a chemical library of 50,000 compounds to identify 93 novel JHR agonists in a reporter system based on *Drosophila* cells. Of these compounds, 26% modulated AHR signaling in an analogous reporter assay in a human cell line, indicating a significant overlap in the agonist repertoires of the two receptors. To explore the structural features of agonist-dependent activation of JHR and AHR, we compared the ligand-binding cavities and their interactions with selective and common ligands of AHR and GCE. Molecular dynamics modeling revealed ligand-specific as well as conserved side chains within the respective cavities. Significance of predicted interactions was supported through site-directed mutagenesis. The results have indicated that synthetic insect juvenile hormone agonists might interfere with AHR signaling in human cells.

## Introduction

Juvenile hormones (JHs) of arthropods comprise a specific group of sesquiterpenoids, absent from vertebrates. In insects, JHs play vital roles by controlling the metamorphosis of larvae to adults^1^ and by stimulating reproduction in the latter.^2^ Like other small and lipophilic signaling molecules, JHs traverse cell boundaries and bind an intracellular receptor protein. Unlike steroids or retinoids, JHs recognize a ligand-dependent transcription factor of the basic helix-loop-helix/PER-ARNT-SIM (bHLH-PAS) family known as methoprene-tolerant (MET).^3–5^ MET was discovered in the fruit fly *Drosophila melanogaster* through screening for mutants surviving exposure to a synthetic JH analog and insecticide methoprene.^6^ The *Drosophila* genome contains a paralogous ancestral gene named *germ cell-expressed* (*gce*), which has duplicated during the evolution of flies.^7^ Genes encoding GCE/MET represent a unique bHLH-PAS protein type that occurs in protostomes but has been lost very early during deuterostomian evolution.^8^ Therefore, vertebrates lack both, the JH and its receptor.

Both of the *Drosophila* paralogous genes encode functional JH receptors (JHRs). GCE and MET bind JH to a cavity within their PAS-B domains,^4,9^ and either protein can substitute for the other in genetic rescue experiments with *Met^-^ gce^-^* double-mutant flies.^9,10^ However, expression of the two JHR genes only partly overlaps *in vivo*^11^ and, therefore, each paralog likely has its specific roles. In some contexts, GCE is the predominantly active JHR protein.^5^ For instance, *Met* mRNA is under-expressed relative to *gce* in a *Drosophila* Kc167 embryonic cell line.^12^ Of relevance to this study, transcriptional activation of JH-inducible reporters in these *Drosophila* cells depends on GCE and its ligand-binding capacity, whereas MET is dispensable for this activation.^9^

To form a transcriptionally active, DNA-binding JHR complex, GCE (or MET) dimerizes with another bHLH-PAS protein called taiman (TAI),^4,13,14^ an insect ortholog of the mammalian nuclear receptor coactivator (NCoA) proteins.^8^ Binding of JH or another agonist to GCE/MET induces its nuclear import,^15^ dissociation from a chaperone complex with HSP90^16,17^ and the ensuing interaction with TAI.^13,14,16–19^ The GCE:TAI or MET:TAI heterodimer binds DNA JH-response elements (JHREs) and initiates transcription of target genes.^13–15,18^ Therefore, the transcriptional activity of the JHR complex is determined by the availability of an agonist ligand.

The above described mode of GCE/MET action closely resembles that of the mammalian aryl hydrocarbon receptor (AHR).^5,8^ Although not orthologous, both proteins share a similar architecture of functional domains including regions bound by HSP90 in the absence of agonists, and regions mediating dimerization with bHLH-PAS partners when agonists bind to the cognate PAS-B cavities.^8,20,21^ One difference is that GCE/MET pairs with TAI, whereas AHR forms a heterodimer with the AHR nuclear translocator (ARNT) that is only remotely related to NCoA.^8^

The most striking common feature of AHR^22–24^ and GCE/MET^9,25,26^ is that both proteins can be activated by chemically diverse, low-molecular weight ligands. AHR agonists range from environmental toxins including polycyclic aromatic hydrocarbons (PAHs) and PAH-like chemicals such as ý-naphthoflavone (β-NF), products of tryptophan degradation (*e.g*., 6-formylindolo[3,2-b]carbazole; FICZ) to metabolites of the intestinal microflora such as indirubin^22–24^. A cryo-EM structure of human AHR bound by indirubin has been recently resolved.^27^ It emerges that the diverse ligands elicit different gene-regulatory activities of AHR.^22^ Given that AHR has been functionally linked with carcinogenesis and immune-related diseases, the search for small molecules that modify AHR activity holds a relevant therapeutic potential. A large-scale high-throughput screening (HTS) has identified thousands of putative chemical agonists and antagonists of the AHR-dependent transcriptional activity.^28^ Initial attempts to identify novel modulators of insect JHR signaling either through chemical HTS based on a cell line from the silkworm, *Bombyx mori*^29,30^ or using a virtual screening approach^31^ have recently been reported.

To our knowledge, GCE/MET presents the first example of a bHLH-PAS protein acting as a hormone receptor. In addition to native insect JHs, the GCE/MET proteins bind a large diversity of synthetic small molecules.^9,25,32^ A few of those, including methoprene, fenoxycarb and pyriproxyfen, have been used as pesticides of the “insect growth regulator (IGR)” group since the mid-1970s, decades before the discovery of their target molecule.^26^ Our previous work has shown that these JH-mimicking compounds, dubbed juvenoids, are highly potent agonist ligands of the GCE/MET proteins. Given the absence of JH in vertebrates, JH mimics are generally considered safe for humans and livestock. However, the existing commercial juvenoids affect non-insect organisms, particularly aquatic, including crustaceans, vertebrates, or plants.^33–35^ This has raised concerns about negative environmental impact of these hormonal mimics as potential endocrine disruptors.^36^ Besides crustaceans that are close relatives of insects possessing the JHR protein orthologs,^35^ molecular targets of JHR agonists in vertebrates are as yet unidentified. The functional parallel in JHR and AHR mode of action, together with well-recognized effects of AHR on endocrine homeostasis of mammals^37,38^ prompted us to examine potential overlaps in agonists activating these two analogous receptor systems.

In this communication, we attempted structural modeling to find similarities in the ligand-binding domains of *Drosophila* GCE and human AHR. To explore the agonist repertoire of GCE, we performed an unbiased HTS for activators of the JHR complex (GCE:TAI) using JHRE-luciferase reporter assays in a *Drosophila* cell line. Validated JHR agonists were then tested in an AHR-dependent reporter assay in human cells. This approach uncovered a significant fraction of novel JHR agonists that also modulated AHR activity. Molecular modeling illustrated that both receptors engage in van der Waals interactions and specific hydrogen bonds with the hydrophobic portions and polar groups, respectively, of their ligands, often via structurally conserved residues that are involved in ligand recognition.

## Results

### Structural implications for GCE signaling mechanism

Structures of AHR complexes, recently resolved by X-ray crystallography^39,40^ and cryo-EM of human AHR in a chaperone complex with HSP90 and XAP2^27^ provided new insights into bHLH-PAS dimerization and ligand binding, respectively. In the absence of experimentally determined JHR structure, we resorted to computational modeling. Using AlphaFold2^41^, we generated models of GCE bound by the *Drosophila* HSP90 ortholog (HSP83) and of the GCE:TAI heterodimer, and compared them with the reported structure of the cytoplasmic, pre-signaling AHR:HSP90:XAP2 complex.^27^

In the AHR complex, PAS-B and its N-terminal linker to PAS-A are threaded through the HSP90 dimer (Figure 1 (A)). Similarly, association of GCE with HSP83 obscures the PAS-A/B linker that is part of the predicted GCE:TAI heterodimer interface, in which both PAS-A and PAS-B together with their linker form an X-shaped assembly (Figure 1 (B,C)). The C-terminal regions of HSP83 would also sterically interfere with the TAI PAS-A. The models suggest that GCE:TAI may be activated through the same mechanism as proposed for AHR, in which the PAS-B of the ligand-free receptor is blocked by HSP90 until a ligand is bound and the HSP90 ATPase action completed with the release of inorganic phosphate and ADP.^27^ Like AHR, the chaperone-bound GCE will not dimerize with its partner TAI to form the bipartite basic DNA binding domain, similar to that of AHR:ARNT.^39,40^

**Figure 1.**
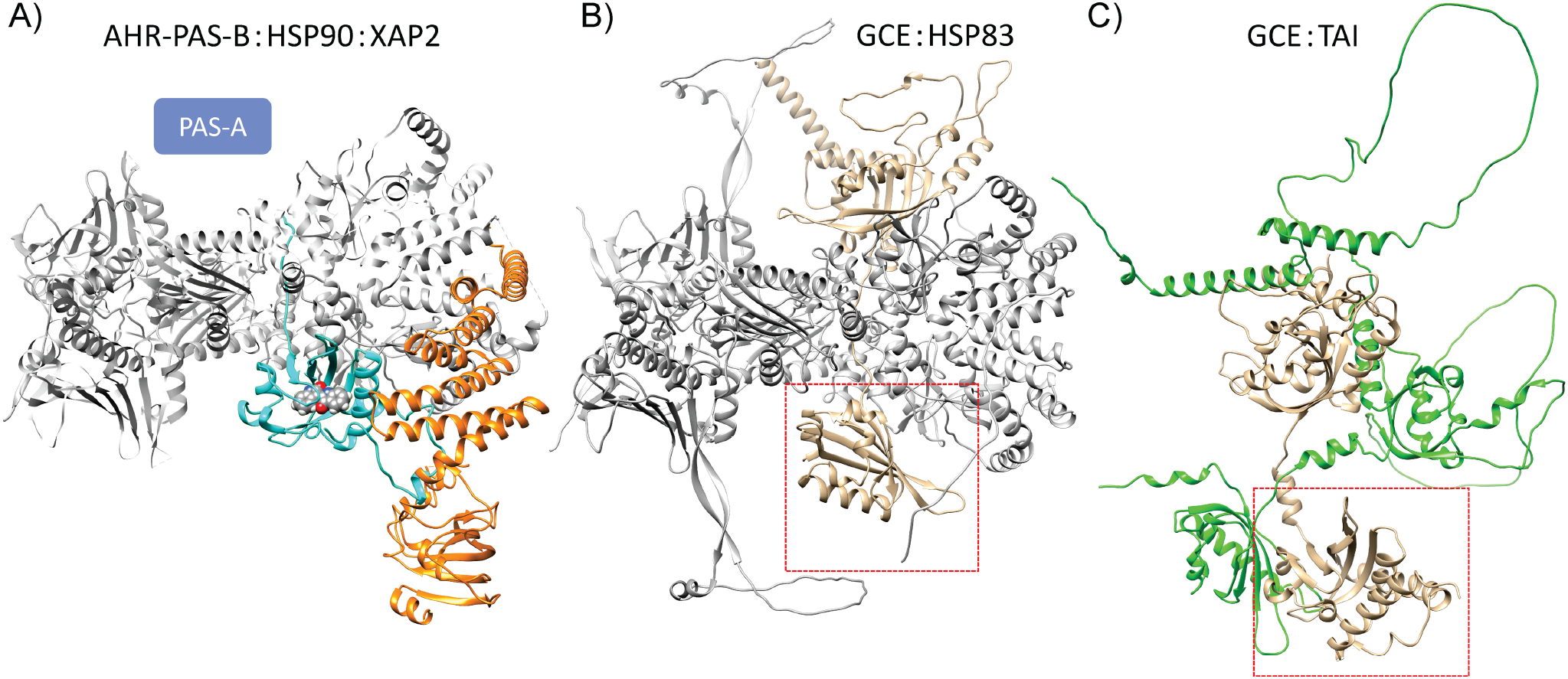
Structural comparison of the human AHR-PAS-B:HSP90:XAP2 complex with models of *Drosophila* complexes of GCE with HSP83 and TAI. A) AHR-PAS-B:HSP90:XAP2 (PDB 7ZUB)^27^ in ribbon representation: cyan, AHR-PAS-B; orange, XAP2; grey, HSP90 dimer. Indirubin is depicted in space fill representation (CPK color) within the PAS-B. The putative position of PAS-A (not resolved in the structure) is indicated. B,C) In the AlphaFold2 models, GCE is colored wheat with the PAS-B outlined in the red boxes. The two HSP83 monomers (B) are depicted in light and darker grey, respectively; TAI is shown in green (C). Structures in panels A and B were orientated using HSP dimer RMSD. The GCE PAS-B domain was used for RMSD alignment of GCE:TAI (C) with the AHR complex (A).

### Similarity of PAS-B ligand binding domains in GCE and AHR

The structure of the human AHR PAS-B domain^27^ revealed a cavity of 682 Å^3^, partially occupied by the natural agonist indirubin. Among the numerous residues within the ligand-binding pocket, H291, F295, and two residues forming hydrogen bonds (S365 and Q383) make contacts with the ligand (Figure 2 (A,B)). Substituting alanine for H291, F295 or S365 (but not Q383) abolished indirubin binding.^27^ To compare the ligand binding mechanism between AHR and GCE, we took the modeling approach previously employed to predict affinities for synthetic JHR agonists.^32^

**Figure 2.**
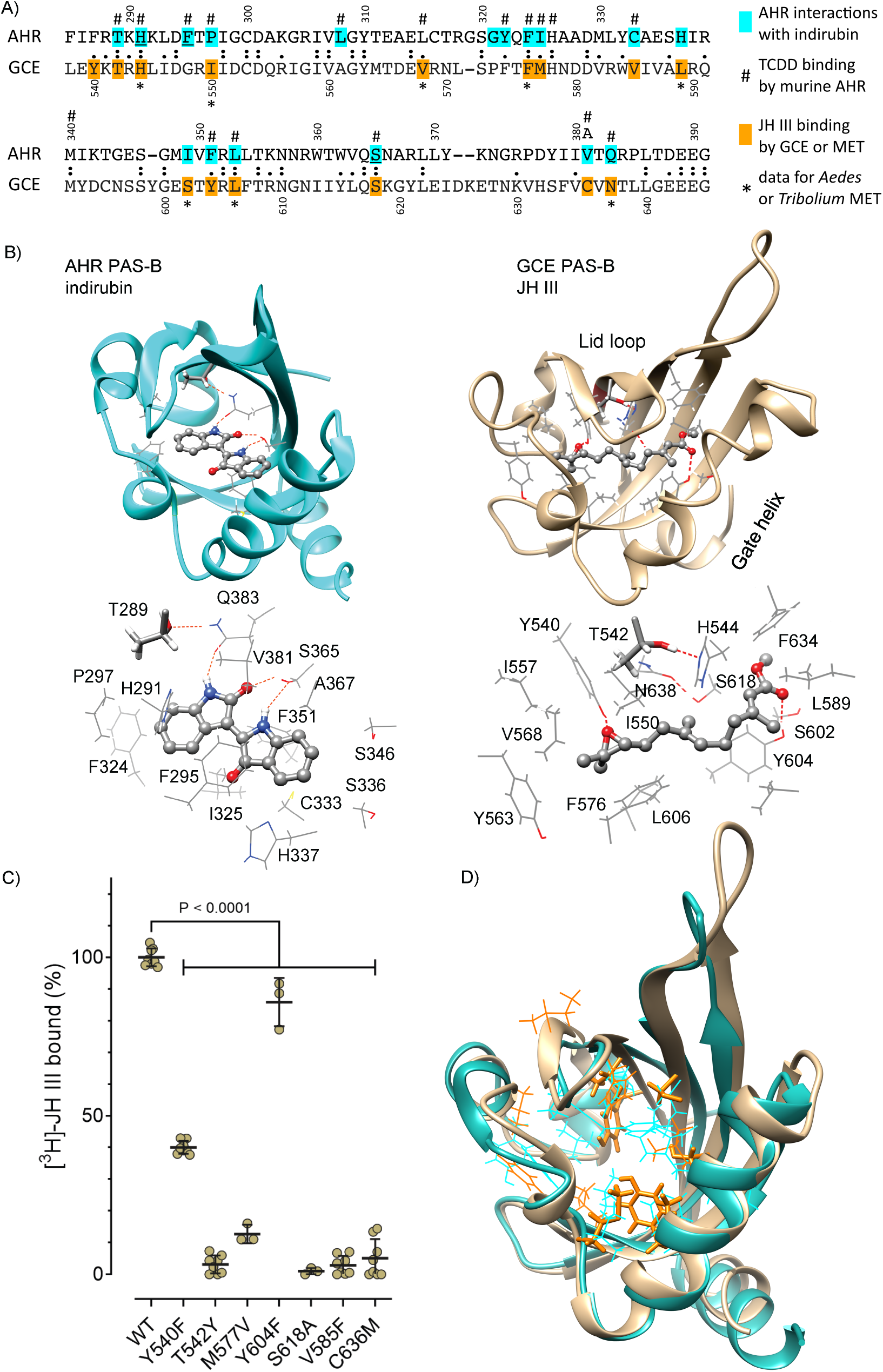
Interactions of AHR and GCE ligand-binding domains with their respective native agonists. A) Alignment of the PAS-B domains of human AHR and GCE. Residues highlighted in cyan engage in diverse interactions of AHR with indirubin; mutating any of the three underlined amino acids prevents indirubin binding.^27^ Hashtags denote residues involved in TCDD binding by murine AHR.^42,43^ The position of V381 in human AHR is occupied by alanine in murine AHR. Residues highlighted in orange are required for binding of JH III by *Drosophila* GCE (C); asterisks indicate JH III binding for conserved residues by *Aedes*^14^ and/or *Tribolium*^4^ GCE/MET orthologs. B) Interactions of PAS-B domains of human AHR (cyan) with indirubin (based on Gruszczyk et al.)^27^ and of GCE (wheat) with JH III. Frames with the highest affinity (as measured by GBSA interaction energy) were selected to reveal the maximum number of potential contacts. The protein backbones are shown in ribbon representation and the key interacting residues (wireframe) with hydrogen bonds are depicted as red dashed lines. The bottom panels identify interacting residues and their hydrogen bonds (red dashed lines). Ligands appear as ball and stick (CPK coloring). The conserved T289/T542 residues in both proteins are highlighted in stick representation. C) Binding of radiolabeled JH III to the wild-type (WT; 100%) and mutated versions of the recombinant GCE protein expressed *in vitro*. Dots represent data from independent assays, shown as mean ± S.D. Differences between each mutated variant and the WT are statistically significant (*P* < 0.0001) based on one-way ANOVA with multiple pairwise comparisons. D) Structural overlay (C_α_ RMSD) of AHR (cyan) and GCE (wheat) ligand binding domains with ligand engaged residues shown in wireframe (colored as highlighted in alignment, A). The GCE residues corresponding to mutations presented in (C) are shown in stick representation.

The GCE PAS-B domain with its native ligand JH III docked inside the cavity was relaxed through a microsecond simulation. A frame with the highest ligand affinity, as judged by the interaction energy (GBSA approximation), was selected to reveal the key contacts and interaction pattern within the binding pocket. Similar to AHR, the GCE cavity is lined by nonpolar and hydrophobic side chains as well as tyrosine and other polar residues (*e.g.*, Y540, T542, H544, S602, Y604, S618, N638) but no charged species (Figure 2 (B)). The hydroxyl groups of Y540 and Y604 engage in hydrogen bonds with the JH III epoxide and ester carbonyl groups, respectively. The significance of both hydrogen bonds is supported by mutating to phenylalanine either Y540^25^ or Y604 (Figure 2 (C)); either mutation partially but significantly reduced binding of JH III to GCE. At the position equivalent to the critical S365 in AHR, mutation of S618 in GCE proved detrimental to JH III binding (Figure 2 (C)). In addition, conserved residues at positions of H544, I550, V568, F576, L606, or N638 in GCE were identified as being important for JH III binding by MET from the mosquito *Aedes aegypti*^14^ or the beetle *Tribolium castaneum*^4^ (Figure 2 (A)).

Interestingly, T542 whose mutation to tyrosine also abolishes the hormone binding (Figure 2 (C)) and transcriptional and *in vivo* activities of GCE^9^, may not contribute to ligand binding directly. The GCE model suggests that T542 engages in a hydrogen bond to H544 from the neighboring β-strand and in stabilizing the cavity (Figure 2 (B)). In the T542Y mutant, this hydrogen bond cannot form, and the bulky Tyr residue takes up substantial volume of the binding cavity, effectively forcing the ligand out towards the “gate” and out of the pocket as illustrated by snapshots from a representative molecular dynamics (MD) trajectory (Figure S1). The ligand seems to exit via the gate which is formed by the gate helix Fα and the lid loop between β-strands A and B.^32^ Another pair of polar residues, the critical S618 (Figure 2 (C)) and N638, also appears to stabilize the GCE cavity by forming a hydrogen bond between the neighboring β-strands (Figure 2 (B)).

In our model of AHR PAS-B, T289 (homologous to T542 in GCE; Figure 2 (A)) forms a hydrogen bond with Q383, which stabilizes the latter in an orientation suitable for binding indirubin (Figure 2 (B)). While experimental data for T289 mutagenesis are unavailable for human AHR, T283M mutation of the equivalent residue within murine AHR (mAHR) has been shown to abolish binding of 2,3,7,8-tetrachlorodibenzo-*p*-dioxin (TCDD).^42,43^ Indeed, Figure S1 shows that *in silico* mutation and MD simulation in a mAHR PAS-B model that was based on a cryo-EM structure (PDB 8H77)^44^ led to the exit of TCDD through the gate, *i.e.*, in a fashion similar to the exit of JH III from the GCE cavity.

Although GCE and AHR are not orthologous, sequence conservation (Figure 2 (A)) and structural overlap of the ligand binding residues (Figure 2 (D)) revealed fundamental analogies between these two receptors. The above data led us to investigate a hypothesis that the observed structural and functional similarities may also be reflected in the ligands, *i.e.*, the chemical structures recognized by these receptors. While thousands of agonists have been reported for AHR,^22,28^ few are known for GCE/MET, preventing a comparative analysis based on existing data. Therefore, we designed and performed an unbiased HTS of a large chemical library for new GCE agonists, followed by testing their activities in a previously established AHR reporter assay.^45^

### High-throughput screening yields diverse agonists of *Drosophila* JHR

We have developed a cell-based reporter system for determination of specific ligand-dependent transactivation of *Drosophila* JHR, amenable to HTS of large chemical libraries. Two reporters, each containing 8 copies of a JHRE placed upstream of a minimal promoter^9,25^ and either the firefly or NanoLuc® luciferase coding sequence (Figure 3 (A)) were stably integrated to the genome of *Drosophila* Kc167 cells. The reporters respond to JHR agonists in a manner dependent on the endogenous GCE and TAI proteins,^9,25^ both of which are naturally expressed in the Kc167 cells.^12^ In contrast, *Met* mRNA is scarce in this cell line,^12^ and MET does not seem to contribute to the JHRE reporter expression.^9,25^ The assay does not respond to agonists when JHRE is mutated (*mut*JHRE)^9^ to prevent binding of the JHR complex, thus providing a specificity control (Figure 3 (B)).

**Figure 3.**
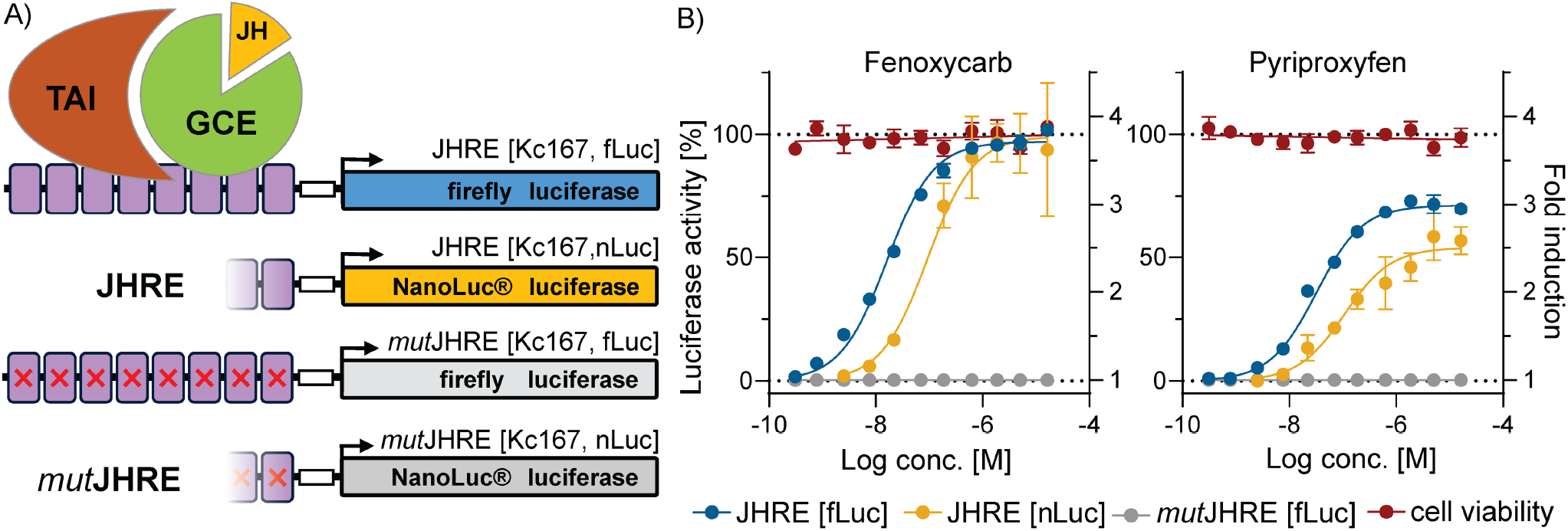
*Drosophila melanogaster* JHR reporter assays. A) A schematic of the luciferase reporter constructs carrying 8 copies of the JHRE (violet boxes) or their mutated version (*mut*JHRE, red crosses) incapable of binding the JHR heterodimer GCE:TAI. The JHREs are linked to a minimal promoter (white box) followed by the coding sequence for either the firefly [fLuc] or NanoLuc® [nLuc] luciferases. The reporter constructs are stably integrated into the genome of the *Drosophila* Kc167 cell line expressing endogenous GCE and TAI. B) A titration experiment with reference compounds fenoxycarb and pyriproxyfen using JHRE [fLuc] (blue) and JHRE [nLuc] (yellow) reporter assays to evaluate JHR activation. The *mut*JHRE [fLuc] reporter (grey) and cell viability (red) assays provide controls for nonspecific and cytotoxic effects. The reporter activities are expressed as percentage of the activities obtained in cells treated with DMSO (0%) and 100 nM fenoxycarb (100%). The *mut*JHRE data are presented as fold induction relative to the basal level of the reporter signal in DMSO-treated cells. Cell viability is indicated relative to cells treated with media alone (0%) and DMSO (100%).

We initially utilized the JHRE [Kc167, fLuc] reporter cell line (Figure 3 (A)) to screen 50,000 diverse drug-like compounds with molecular weights between 150 and 559 in an automated system. The flow chart presenting the screening process is shown in Figure S2 and details can be found in Materials and Methods. Each compound was tested in duplicate at a single concentration of 10 μM. The data were processed and normalized using the Bscore algorithm^46^ to compensate for potential artifacts introduced by the instrumentation (Figure 4 (A)). Compounds with a Bscore ≥ 5 were then retested in dose-response experiments with the same reporter assay and with the *mut*JHRE reporter assay. This step eliminated false positives and nonspecific activators of the reporter assay. In the next step, compounds with confirmed activity were retested in the JHRE [nLuc] assay where the NanoLuc® luciferase replaces fLuc to eliminate nonspecific activators of the firefly luciferase itself. The remaining compounds were tested in a cell viability assay to determine potential cytotoxicity of the compounds (Figure 3 (B)). As a result, we identified and validated 93 new agonists of the *Drosophila* JHR with the average potency in the firefly and NanoLuc® luciferase reporter assays ranging from 600 nM to 70 μM (Figure 4 (B)). None of the novel agonists exceeded the potency of the reference compounds fenoxycarb (EC_50_ = 58 nM) or pyriproxyfen (EC_50_ = 76 nM). Examples of four individual agonists eliciting distinct types of specific JHR response (compounds 1.1 through 1.4) are shown in Figure 4 (C) and activities of all discovered agonists are summarized in Table S1.

**Figure 4.**
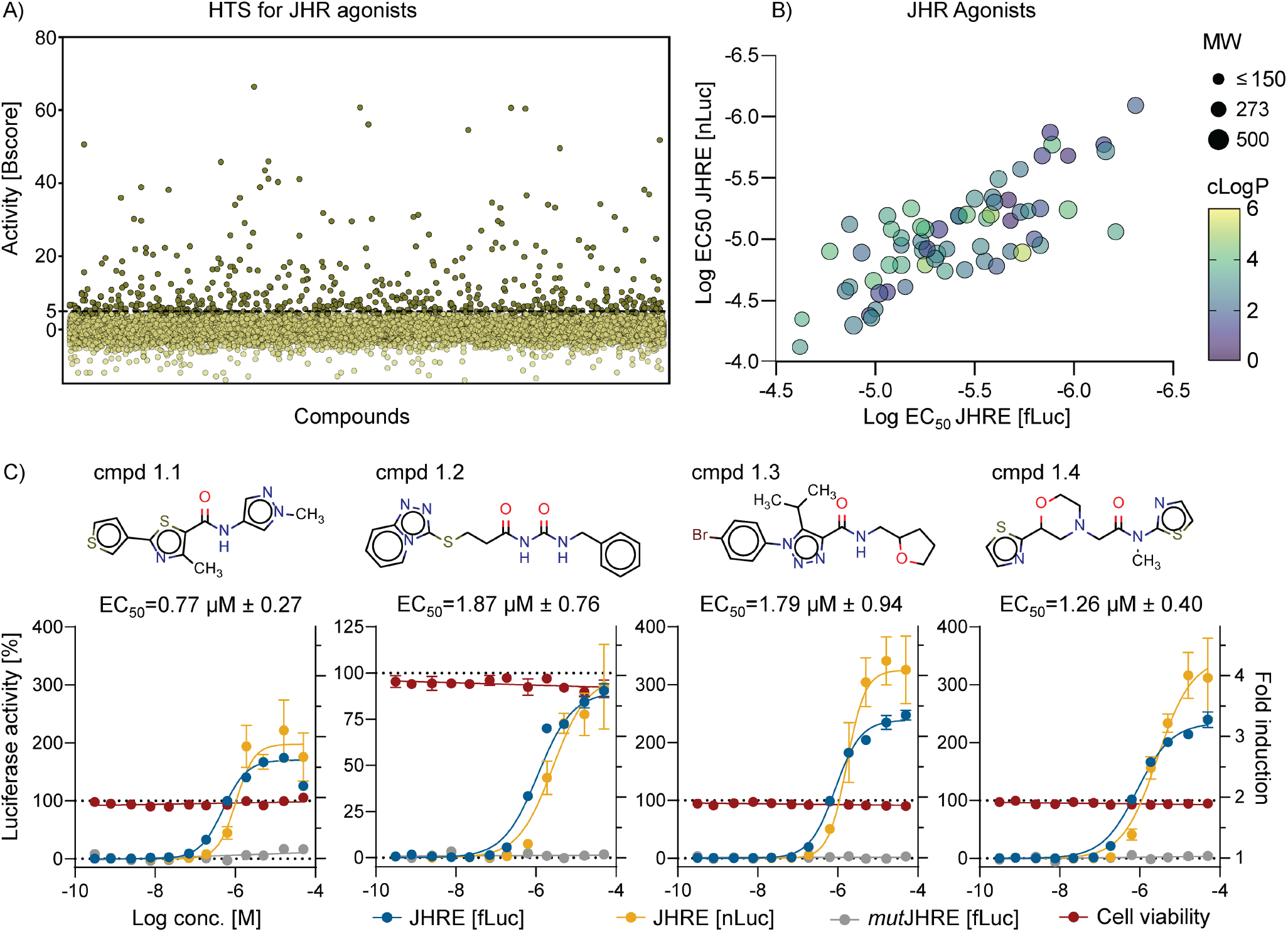
High-throughput screening (HTS) of a compound library for novel *Drosophila* JHR agonists. A) Activities of 50,000 individual compounds screened with the JHRE [fLuc] reporter assay at 10 μM concentration in duplicates. Compounds with the Bscore ≥ 5 were selected as hits. B) Correlation of potencies (EC_50_) of validated JHR agonists in the JHRE [fLuc] and JHRE [nLuc] reporter assays. The dot size represents molecular weight of each compound and lipophilicity (cLogP) is indicated by the color scale from cyan to yellow. C) Examples of chemical structures and potencies of validated JHR agonists as tested in dose response experiments using the JHRE [fLuc] (blue), JHRE [nLuc] (yellow), *mut*JHRE [fLuc] (grey), and the cell viability (red) assays. Data are expressed as percentage of the activities in cells treated with DMSO (0%) and 100 nM fenoxycarb (100%). In the *mut*JHRE assay, activities are presented as fold induction relative to the basal reporter signal in DMSO-treated cells. Cell viability is indicated relative to cells treated with medium alone (0%) and DMSO (100%). EC_50_ data are reported as mean ± S.D. of EC_50_ values obtained from JHRE [fLuc] and JHRE [nLuc] experiments.

### Interactions of GCE and AHR with selective agonists

Given the conserved features of the PAS-B domains of GCE and AHR (Figure 2 (A,B,D)), we investigated to what extent agonists of JHR and AHR, respectively, might exert activity towards the analogous but evolutionarily distant human and *Drosophila* receptors. To achieve this, we employed a clonal luciferase reporter cell line AZ-AHR described by Novotna et al.,^45^ which responds to the transcriptionally activated AHR. This reporter is driven by multiple copies of the dioxin response elements (DRE) recognized by AHR. The reporter was stably integrated in the genome of the human hepatocellular carcinoma cell line HepG2.^45^

First, we assembled a collection of established AHR agonists: indirubin, benzo[a]pyrene (B[a]P) and FICZ; juvenile hormones: JH III (the most common natural analog), JH III bisepoxide (JHB_3_, specific to flies), methyl farnesoate (MF) and farnesol (both precursors in JH biosynthesis); and a set of commercial juvenoids: fenoxycarb, pyriproxyfen, S-methoprene, and S-hydroprene. Each compound was tested at a single concentration in the *Drosophila* JHR and human AHR reporter assays (Figure 5 (A,B)). In the next step, potency of the active compounds was characterized in a wide range of concentrations (Figure 5 (C,D)). As expected, both the natural hormones and the juvenoids showed agonistic activity in the JHR reporter assay while the AHR agonists activated the AHR reporter (Figure 5 (A,B)). We found a marginal AHR activation for pyriproxyfen with only 7 ± 2% efficacy compared to 100% attained with 25 μM B[a]P (Figure 5 (B)). Similarly, FICZ weakly activated the JHR reporter exhibiting 7 ± 6% efficacy relative to the activity of 100 nM fenoxycarb (Figure 5 (A)). These data indicate that well-established ligands of both receptors are selective agonists to each of them.

**Figure 5.**
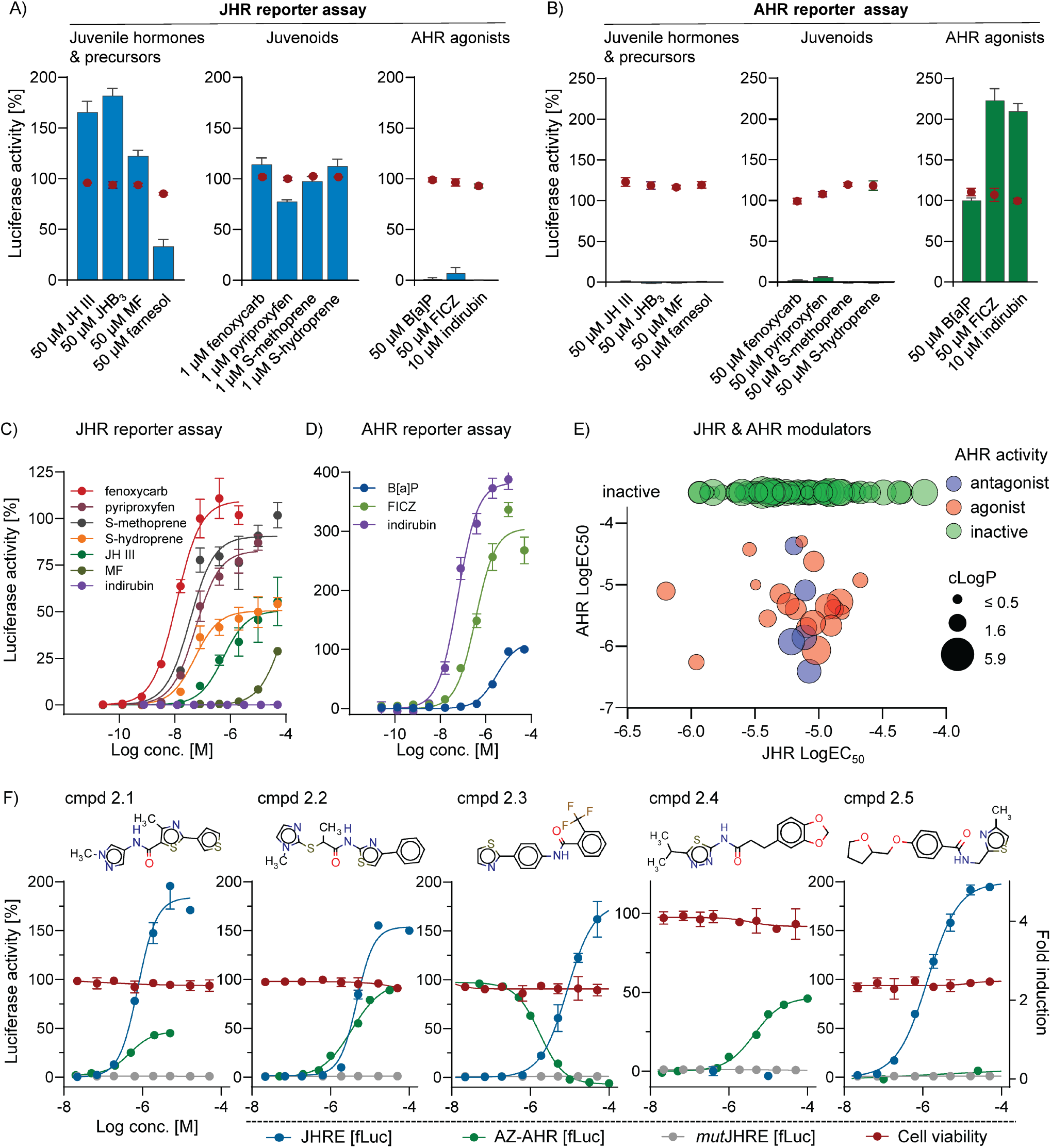
Common and unique agonists of JHR and AHR and their activities in reporter assays. A,B) Juvenile hormones, their precursors and synthetic juvenoids activate specifically JHR while AHR ligands are specific for AHR. Activities are expressed relative to 100 nM fenoxycarb and 25 μM B[a]P representing 100% of activity in the JHR and AHR reporter assays, respectively. Cell viability was assessed in a parallel experiment and activities are represented by red dots. C,D) Titration experiments with specific JHR and AHR agonists. E) Potencies (EC_50_ values) of 93 novel JHR agonists as determined in both AHR [fLuc] and JHRE [fLuc] reporter assays. Color code distinguishes between activities towards AHR: agonist (red), antagonist (blue), inactive (green). The dot size is proportional to the calculated lipophilicity of each compound. F) Examples of chemical structures and responses to the individual compounds active as agonists common to JHR and AHR (cmpds 2.1 and 2.2), a JHR agonist and AHR antagonist (cmpd 2.3), and selective agonists of either AHR (cmpd 2.4) or JHR (cmpd 2.5), respectively. Compounds were tested in dose-response experiments in the JHRE [fLuc] (blue), AZ-AHR [fLuc] (green), and *mut*JHRE [fLuc] (grey) reporter assays and for cell viability (red). The data are expressed relative to activities of 100 nM fenoxycarb and 25 μM B[a]P, which are set to 100% in the JHRE and AZ-AHR assays, respectively. Data for the *mut*JHRE and cell viability assays are shown as in Figure 4 (C).

Structurally, the JHR selectivity for fenoxycarb can be explained by the cavity shape, distinct from that of AHR. Putative binding modes for fenoxycarb docked to GCE and AHR are compared after MD equilibration in Figure S3. The elongated fenoxycarb fit well into the GCE pocket, which became only slightly enlarged (498 Å^3^) relative to being bound by JH III (446 Å^3^). As with JH III, Y540 formed a hydrogen bond with fenoxycarb. However, compared with JH III, fenoxycarb engaged more (19 *vs*. 14) residues in the cavity (Figure S3), which may explain its superior agonist potency and binding affinity to GCE.^25^ In contrast, after being docked in place of indirubin to the AHR PAS-B cavity and equilibrated for at least 1 μs, fenoxycarb partially escaped to a secondary binding pocket (also identified in the cryo-EM structure^27^). Fenoxycarb made 14 contacts and no hydrogen bond with the AHR cavity (Figure S3). This relatively weak and non-specific engagement at a site close to the gate, which was identified as the ligand exit gateway for the mutant variants of both receptors (Figure S1), might represent a metastable state eventually leading to ligand release, although this was not observed during our simulations on a microsecond time scale.

### Common and unique agonists of JHR and AHR

To broaden the studied chemical diversity, we next tested the 93 novel JHR agonists uncovered in the HTS for AHR activity using the AZ-AHR reporter assay. First, we pre-tested all compounds at two doses of 100 nM and 25 μM to identify active molecules. Ligands can elicit either agonist or antagonist activities upon binding to a receptor. Therefore, we carried out an independent experiment with the canonical AHR ligand TCDD, added at a 10 nM concentration to reveal potential antagonists along with agonists. Compounds active in either type of assay were then tested in a wide range of concentrations and EC_50_ values were calculated for each compound. A compound with an EC_50_ below 50 μM was recognized as an AHR agonist or antagonist.

The results showed a surprising overlap of activities. In total, we found 24 compounds that modulated human AHR activity in the set of 93 agonists of *Drosophila* JHR, representing a 26% overlap. Of these, 19 were agonists and 5 were antagonists of AHR (Figure 5 (E)). Examples of different structures of compounds and their distinct effects on AHR activity are shown in Figure 5 (F). Compounds 2.1 and 2.2 exemplify dual agonists of both JHR and AHR, whereas compound 2.3 is at the same time a JHR agonist and AHR antagonist. Compounds 2.4 and 2.5 display selective agonism towards either of the two receptors (Figure 5 (F)).

To gain insight into the dual agonist mode, we docked compound 2.1 into GCE and AHR PAS-B in place of their native ligands, JH III and indirubin, respectively. After microsecond-long equilibration of each MD ensemble (sufficient to reach a stationary state), we compared two representative frames with highest affinity based on GBSA interaction energy approximation from each simulation (Figure S4). Compound 2.1 fit the elongated GCE cavity in a fashion similar to JH III but, being shorter, engaged fewer residues (compare Figure 2 (B) and Figure S4 (A)), some of which overlap with the native ligand contacts (T542, H544, V585, Y604) while also recruiting other side chains. The internal cavity was enlarged to 770 Å^3^ relative to JH III-bound state, reflecting a mismatch between the ligand shape and malleability of the receptor structure. The hydrogen bond between the conserved T542 and H544 was present.

In contrast, the orientation of compound 2.1 was almost perpendicular to the AHR PAS-B cavity whose volume (529 Å^3^) was only slightly enlarged relative to the indirubin-bound state, allowing for 10 contacts (of which T289, L308, F324, L353 and Q383 are common with the indirubin containing structure) and extensive aromatic residue stacking around the ligand (Figures 2 (A) and S4 (A)). This is consistent with the slightly higher potency of compound 2.1 towards AHR *vs*. JHR (Table 1). The hydrogen bond between the conserved T289 and Q383 was maintained.

**Table 1.**
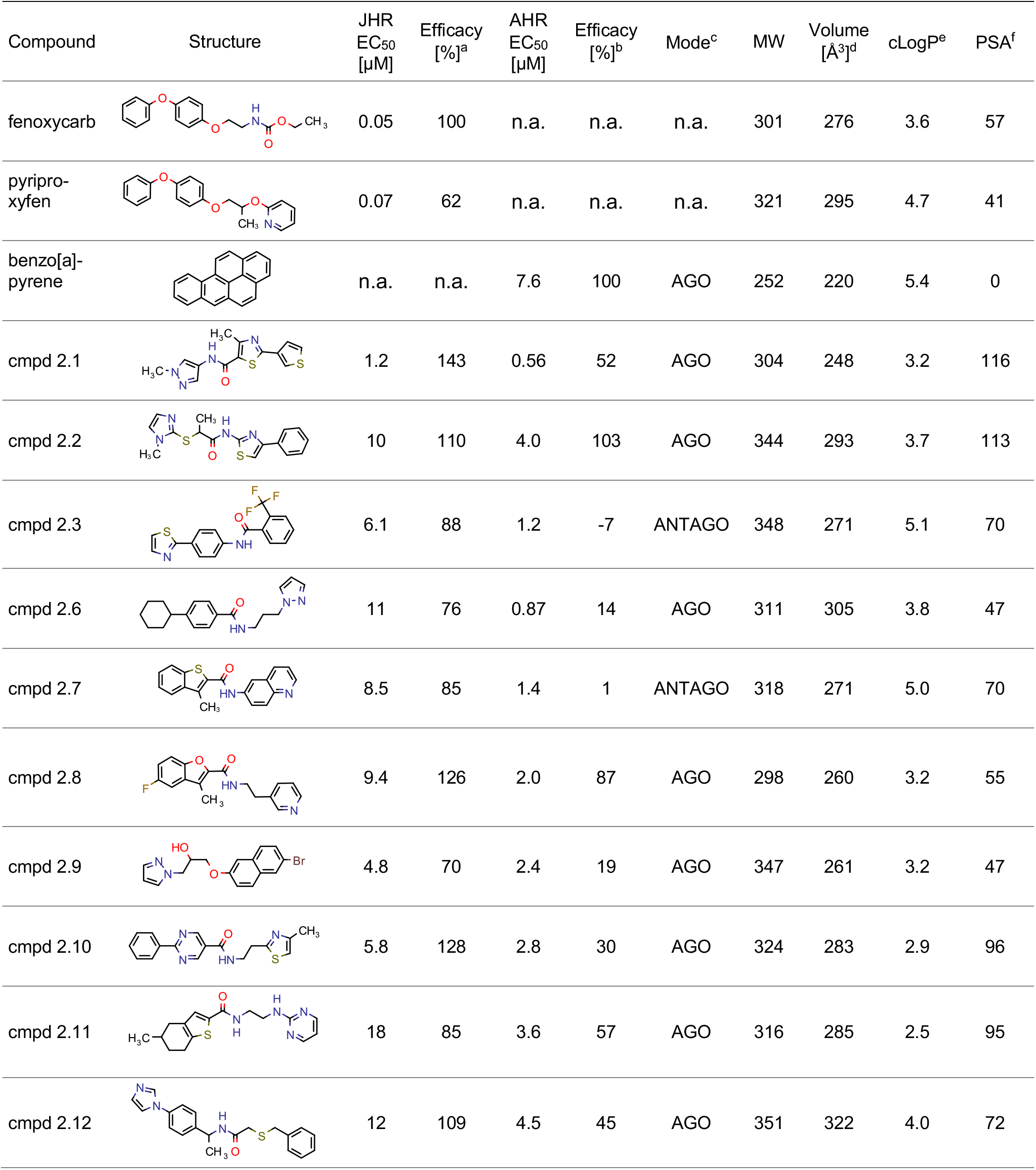
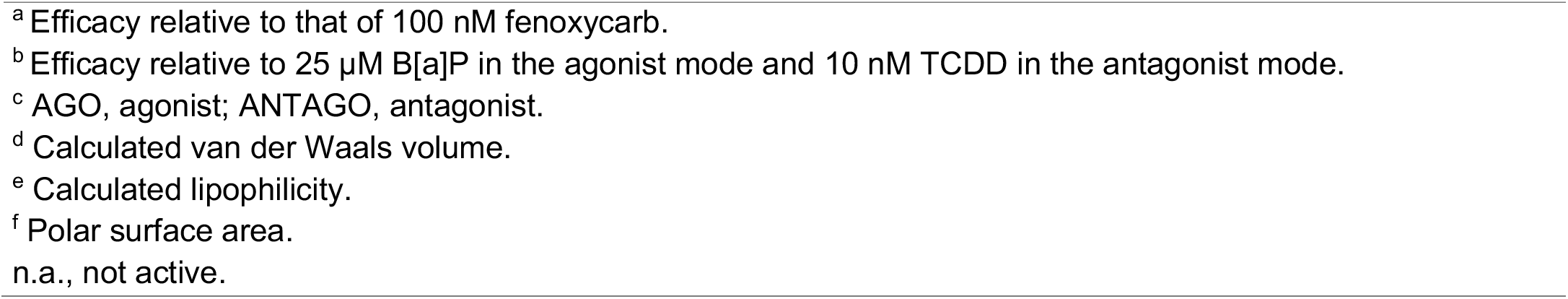
Selected reference agonists and most potent common modulators of JHR and AHR.

| Compound | Structure | JHR<br>EC<sub>50</sub><br>[μM] | Efficacy<br>[%]<sup>a</sup> | AHR<br>EC<sub>50</sub><br>[μM] | Efficacy<br>[%]<sup>b</sup> | Mode<sup>c</sup> | MW | Volume<br>[Å<sup>3</sup>]<sup>d</sup> | cLogP<sup>e</sup> | PSA<sup>f</sup> |
| --- | --- | --- | --- | --- | --- | --- | --- | --- | --- | --- |
| fenoxycarb |  | 0.05 | 100 | n.a. | n.a. | n.a. | 301 | 276 | 3.6 | 57 |
| pyripro-<br>xyfen |  | 0.07 | 62 | n.a. | n.a. | n.a. | 321 | 295 | 4.7 | 41 |
| benzo[a]-<br>pyrene |  | n.a. | n.a. | 7.6 | 100 | AGO | 252 | 220 | 5.4 | 0 |
| cmpd 2.1 |  | 1.2 | 143 | 0.56 | 52 | AGO | 304 | 248 | 3.2 | 116 |
| cmpd 2.2 |  | 10 | 110 | 4.0 | 103 | AGO | 344 | 293 | 3.7 | 113 |
| cmpd 2.3 |  | 6.1 | 88 | 1.2 | -7 | ANTAGO | 348 | 271 | 5.1 | 70 |
| cmpd 2.6 |  | 11 | 76 | 0.87 | 14 | AGO | 311 | 305 | 3.8 | 47 |
| cmpd 2.7 |  | 8.5 | 85 | 1.4 | 1 | ANTAGO | 318 | 271 | 5.0 | 70 |
| cmpd 2.8 |  | 9.4 | 126 | 2.0 | 87 | AGO | 298 | 260 | 3.2 | 55 |
| cmpd 2.9 |  | 4.8 | 70 | 2.4 | 19 | AGO | 347 | 261 | 3.2 | 47 |
| cmpd 2.10 |  | 5.8 | 128 | 2.8 | 30 | AGO | 324 | 283 | 2.9 | 96 |
| cmpd 2.11 |  | 18 | 85 | 3.6 | 57 | AGO | 316 | 285 | 2.5 | 95 |
| cmpd 2.12 |  | 12 | 109 | 4.5 | 45 | AGO | 351 | 322 | 4.0 | 72 |

<sup>a</sup> Efficacy relative to that of 100 nM fenoxycarb.
<sup>b</sup> Efficacy relative to 25 μM B[a]P in the agonist mode and 10 nM TCDD in the antagonist mode.
<sup>c</sup> AGO, agonist; ANTAGO, antagonist.
<sup>d</sup> Calculated van der Waals volume.
<sup>e</sup> Calculated lipophilicity.
<sup>f</sup> Polar surface area.
n.a., not active.

Interestingly, the slightly larger but similarly elongated compound 2.2 revealed similar orientation after docking and MD equilibration into both receptors (Figure S4 (B)). Their internal cavities became significantly enlarged to 723 Ȧ^3^ and 833 Ȧ^3^ for AHR and GCE, respectively. Similarly, this compound engages with many residues that also engage the native ligands (compare Figure S4 (B) with Figure 2 (B)). As in the case of compound 2.1, GCE binding exhibited fewer contacts than AHR, reflecting their EC_50_ values (Table 1).

Structural diversity of the common modulators of JHR and AHR is illustrated in Table 1, presenting the respective reporter assay activities and physico-chemical properties of the compounds. Interestingly, all of them are more potent on AHR than on JHR although they were initially identified in the screen for JHR agonists. The most prominent example is compound 2.6, being 13-fold more potent agonist of AHR than of JHR (Table 1).

Physico-chemical analysis provides additional insights into the chemical character of JHR/AHR modulators. In addition to our data, we examined a dataset from an independent HTS for AHR agonists reported in PubChem as AID2796.^28^ The authors used a similar reporter gene assay to the one used in this study, with HepG2 cells stably transfected with a reporter construct carrying a dioxin-responsive promoter.^47^ Their primary screening of 324,747 compounds yielded 7,988 agonists active at 5 μM. Our present analysis of the data revealed a shift towards lower molecular weight in the compounds active on both JHR and AHR relative to the distribution of molecular weights in the screening library (Figure 6 (A)). Specifically, the molecular weight distribution in the screening library ranged from 150 to 559, with the highest frequency occurring at 351. The distribution for JHR agonists peaked at 329, and for AHR modulators at 321 (Figure 6 (A)). The heaviest JHR and AHR agonists had molecular weights of 398 and 381, respectively. A similar shift in the distribution profile was observed for AHR agonists in the PubChem AID2796^28^ which was not biased by a preselection with JHR agonists. The distribution of molecular weights reached a maximum at 323 for AHR agonists, compared to 358 for the whole library (Figure 6 (B)).

**Figure 6.**
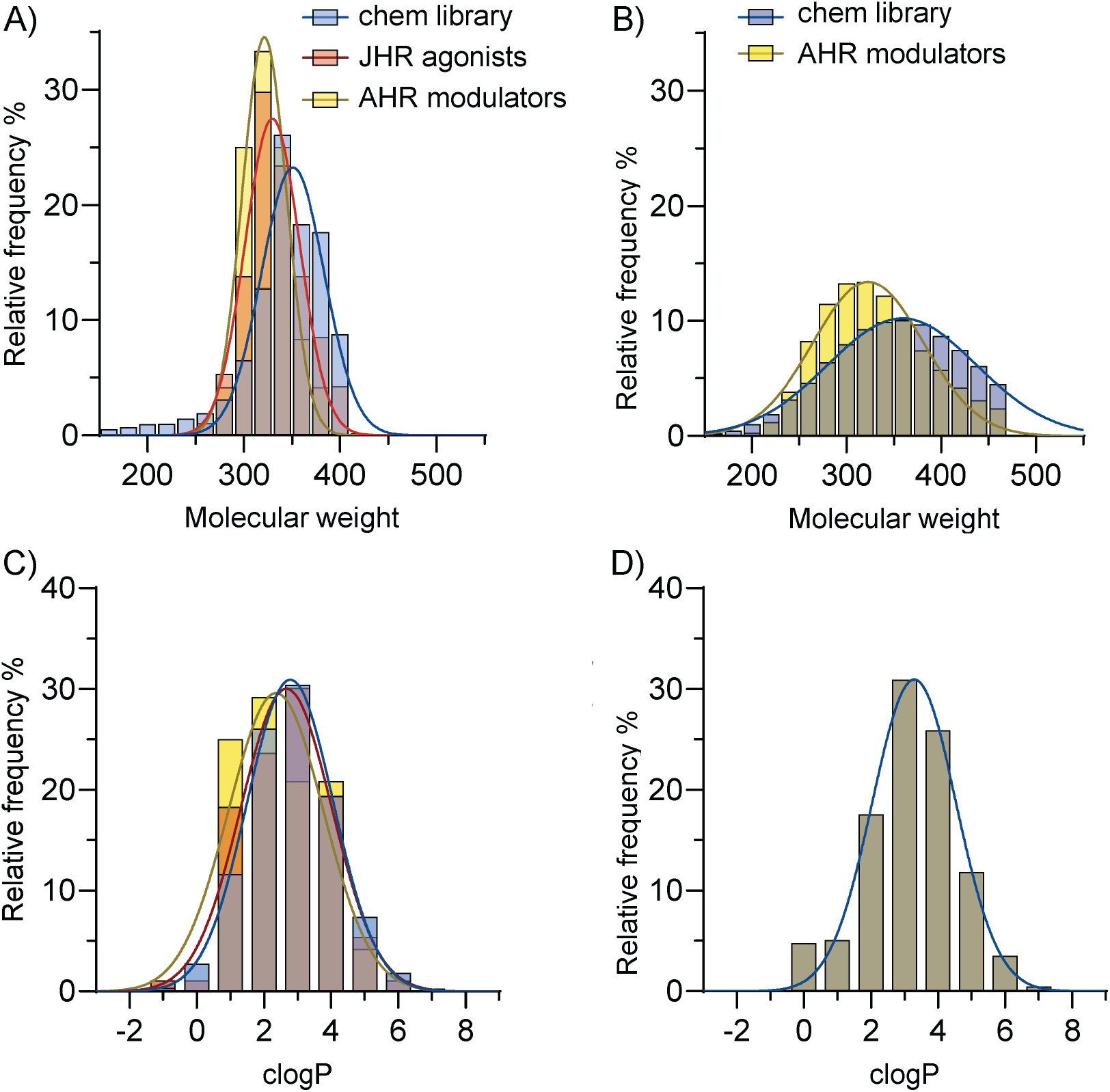
Physico-chemical properties of compound libraries and of JHR and AHR modulators. Distributions of molecular weight (A,B) and lipophilicity (C,D) of compounds present in the current compound library (A,C) and in the HTS PubChem AID2796^28^ (B,D). Profiles of the screening libraries (blue bars) are compared to JHR agonists (orange bars) and AHR modulators (yellow bars) found in the respective libraries.

No apparent trend emerged for lipophilicity of the new modulators (Figure 6 (C,D)). Most of them show lipophilicity between 2.5 and 3.5, *i.e.*, properties which drug-like chemical libraries are optimized for. However, reference ligands of both receptors tend to be more lipophilic: pyriproxyfen (cLogP = 4.7), fenoxycarb (cLogP = 3.6) (Table 1), and B[a]P (cLogP = 5.4). Among the common agonists, we found no purely non-polar compounds, in contrast to some selective AHR agonists, *e.g.*, B[a]P and TCDD, which have low polar surface area (PSA) of 0 and 18, respectively. However, the PSA of the common JHR/AHR agonists varied between 47 and 166 (Table 1), thus reflecting the overall more polar character of typical JHR ligands pyriproxyfen and fenoxycarb with PSA of 47 and 57, respectively. In general, most of the discovered new JHR agonists are linear, elongated molecules with no branching in the chain. They contain up to four aromatic (hetero)cycles with occasional aliphatic cycles. Of note, planar groups composed of two aromatic rings are frequently present, such as quinoline and benzothiophene in compound 2.7, benzofurane in compound 2.8, or naphthalene in compound 2.9. Moreover, the rigidity of these molecules is often strengthened with peptidic bonds connecting the rings, as exemplified by compounds 2.3 or 2.7 (Table 1). This observation is further supported by the examples of compound 2.7 and other JHR agonists containing the central thiophene-2-carboxamide motif (Table 2). JHR agonists with a peptide bond connecting two different heterocycles show micromolar agonist activity on AHR. However, increasing of the linker length leads to the loss of activity on AHR.

**Table 2.**
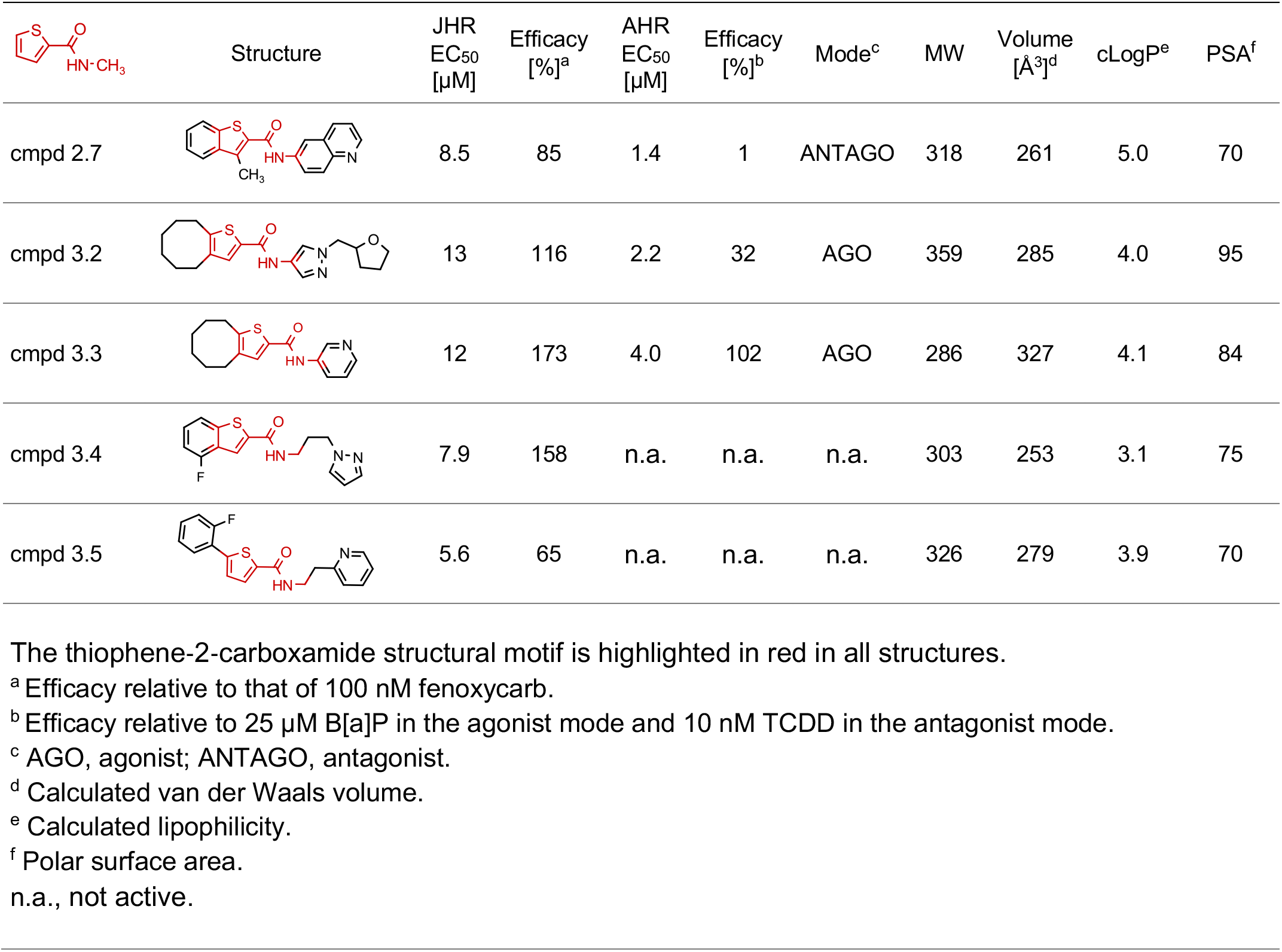
Activities of compounds containing the thiophene-2-carboxamide motif.

| | | Structure | JHR<br>EC<sub>50</sub><br>[μM] | Efficacy<br>[%]<sup>a</sup> | AHR<br>EC<sub>50</sub><br>[μM] | Efficacy<br>[%]<sup>b</sup> | Mode<sup>c</sup> | MW | Volume<br>[$\text{\AA}^3$]<sup>d</sup> | cLogP<sup>e</sup> | PSA<sup>f</sup> |
| --- | --- | --- | --- | --- | --- | --- | --- | --- | --- | --- | --- |
| cmpd 2.7 |  |  | 8.5 | 85 | 1.4 | 1 | ANTAGO | 318 | 261 | 5.0 | 70 |
| cmpd 3.2 |  |  | 13 | 116 | 2.2 | 32 | AGO | 359 | 285 | 4.0 | 95 |
| cmpd 3.3 |  |  | 12 | 173 | 4.0 | 102 | AGO | 286 | 327 | 4.1 | 84 |
| cmpd 3.4 |  |  | 7.9 | 158 | n.a. | n.a. | n.a. | 303 | 253 | 3.1 | 75 |
| cmpd 3.5 |  |  | 5.6 | 65 | n.a. | n.a. | n.a. | 326 | 279 | 3.9 | 70 |

The thiophene-2-carboxamide structural motif is highlighted in red in all structures.
<sup>a</sup> Efficacy relative to that of 100 nM fenoxycarb.
<sup>b</sup> Efficacy relative to 25 μM B[a]P in the agonist mode and 10 nM TCDD in the antagonist mode.
<sup>c</sup> AGO, agonist; ANTAGO, antagonist.
<sup>d</sup> Calculated van der Waals volume.
<sup>e</sup> Calculated lipophilicity.
<sup>f</sup> Polar surface area.
n.a., not active.

### Agonist-induced changes of receptor cavities

To explore how the volume of the receptor cavities responds to different ligand sizes and whether the pockets remain stable in the absence of ligands, we conducted MD simulations of the PAS-B domains with a variety of ligands docked and with ligands removed. Due to the large but random fluctuations in the cavity size, we computed the average cavity volume over the trajectory stationary phase for each state and compared it with the ligand mass (Figure S5). The average cavity volume (356 Ȧ^3^) within the human AHR PAS-B (PDB 7ZUB^27^) remained similar to the ligand filled state (382 Ȧ^3^). This lack of response was confirmed by running a simulation of the empty structure from the murine AHR (PDB 8H77,^44^ cavity 380 Ȧ^3^). In contrast, when emptied of JH III, the GCE PAS-B model reduced its average cavity volume from 522 to 290 Ȧ^3^. As expected, mutations replacing the conserved threonine with a bulkier residue tended to reduce the cavity volume. While both AHR and GCE can expand their cavities in response to larger ligands, a significant trend is only discernible for GCE (linear regression lines in Figure S5).

As the receptor cavities adjust dynamically to the size and shape of ligands, it is difficult to define the volume available for any given ligand. Still, surprisingly to us, the calculated van der Waals volumes of the newly discovered modulators only ranged from 250 to 327 Å^3^ (Table 1), thus falling bellow the theoretically available volume of the AHR cavity.^27^ One reason might be that the chemical library does not contain molecular structures ideally suited to fit the larger cavity space and its specific shape, or such molecules might be sterically hindered to enter the cavity.

## Discussion

Our computational model of *Drosophila* GCE compared to the cryo-EM structure of the human AHR revealed striking similarities between the two receptors. The AlphaFold model suggested that GCE is stabilized in its cytosolic unliganded state by complexing with chaperones in a way similar to AHR and that, upon binding a ligand, GCE is activated by an analogous mechanism. This mode of action is supported by previous studies on the JHR proteins.^16,17,25,32^ While GCE and AHR each recognize own specific ligands that differ in shape, modeling of their PAS-B domains unveiled common features of the binding cavities: (1) largely hydrophobic lining; (2) absence of charged residues; and (3) presence of hydrogen bonds, both between side chains within the receptor cavity and with the ligand. As shown for AHR,^27^ hydrogen bonding to the ligand is important for ligand specificity, and this is true for the GCE:JH III complex as well. The significance of some of the hydrogen-bond forming residues for JH binding to GCE was confirmed through binding assays with radiolabeled JH III previously^25^ and in this study (Figure 2 (C)). Our models of the PAS-B domains revealed partly conserved residues that appear to guide the ligand preferences of the two receptors (Figure 2 and Supplementary material).

Structural similarities between the PAS-B domains of GCE and AHR led us to hypothesize that both receptors might share common ligands. GCE binds sesquiterpene molecules, the juvenile hormones, and artificial juvenoids mimicking the hormonal activity. Hundreds of juvenoids were synthesized over the past 50 years during development of a new generation of insecticides classified insect growth regulators (IGRs), aka insect growth disruptors (IGDs).^48,49^ However, evidence that these molecules interfere with the insect hormonal signaling by activating JHR via binding the GCE or MET proteins is thus far only available for three commercial insecticides: S-methoprene, pyriproxyfen, and fenoxycarb.^4,9,25,26^

The structural diversity of AHR ligands is incomparably higher relative to JHR, as AHR has proven an extremely promiscuous binder. However, no significant cross-activity was observed for canonical reference agonists of either JHR or AHR on the other receptor in this study, and no evidence for common ligands has been reported to date, suggesting that well-established ligands are highly specific and selective. However, when tested against JHR agonists that we have newly identified in a compound library through an unbiased HTS, as many as 26% of validated JHR agonists modulated transcriptional response to AHR.

AHR mediates detoxification, physiological and developmental responses to a broad range of agonists from environmental pollutants to endogenous metabolites to products of microbiota. A class of physiologically relevant AHR ligands are indoles, occurring as either tryptophan and its endogenous metabolites tryptamine and indole-3-acetate,^50^ metabolites produced by the intestinal commensal microbiota such as indirubin, indole-3-aldehyde,^51^ 3-methylindole or indoxyl-3-sulfate.^52^ Diverse potent indole-containing AHR ligands were found in brassica vegetables, such as indole[3,2-b]carbazole, 2-(indol-3-yl methyl)-3,3′-diindolylmethane^53^ or indole-3-acetonitrile. The antimigraine drug avitriptan that contains an indole group is another AHR agonist.^54^ The indole is a frequently occurring motif, present in a total of 1055 compounds in the screening chemical library. None of these compounds were active on JHR, so it is likely that screening initially for JHR agonists excluded potential AHR ligands, and that indole-containing compounds do not fit the GCE PAS-B pocket. Similarly, compounds with multiple fused rings typical for AHR ligands FICZ, TEACOPs, TCDD, B[a]P, did not emerge as common ligands, even though the chemical library included such structures. Therefore, ligands common to GCE and AHR occupy a relatively confined region in the chemical space, defined by a narrow overlap between a larger region comprising AHR ligands and a relatively smaller region of JHR agonists.

Development of juvenoid insecticides started by the late 1960s at the Zoëcon company aiming to produce environmentally safe agents with a hormone disruptive mode of action, targeting developmental processes normally regulated by JH.^48^ Methoprene was the first juvenoid approved for commercial use in 1975 and was soon followed by fenoxycarb and by the currently most universally deployed juvenoid pyriproxyfen introduced in 1996.

There is increasing evidence that many exogenous compounds interfere with the endocrine system by disrupting hormonal regulation with consequences on animal and human health and reproduction. Such chemicals are classified as endocrine disruptive compounds (EDCs) and act through diverse molecular mechanisms.^55^ Since juvenoids directly target insect hormonal regulation, these insecticides are EDCs by nature and may cause environmental harm by affecting diverse non-target invertebrate species to which JH signaling is vital.^33–36^ On the other hand, juvenoids are potentially beneficial as they, in theory, should not affect vertebrates that evolutionarily lack the MET/GCE JHR proteins,^8^ the primary target of both native JHs and juvenoids.^26^

In vertebrates including humans, ligand-binding transcription factors of the nuclear receptor (NR) family and AHR are the most prominent targets for EDCs. Both protein types regulate hormonal homeostasis and mutually interact at multiple levels including direct binding of AHR to NRs such as the estrogen receptor (ER) or the NR coactivator NCoA.^21,56^ In addition to non-peptidic hormones and other signaling molecules including steroids, thyroid hormones, fatty acids, retinoids and vitamin D, the NRs bind a variety of EDCs.^57,58^

HTS studies based on the 10K compound library, collected for the “Toxicology in the 21st Century” screening program (Tox21, https://tox21.gov/)^59^ enabled profiling of 1425 pesticides.^60^ Searching of the databases has revealed that fenoxycarb interacts with a number of NRs such as androgen (AR), estrogen (ERα, ERβ), progesterone (PR), peroxisome proliferator-activated receptor (PPARγ), or thyroid hormone (TRα) receptors. Pyriproxyfen shows fewer interactions, including AR, ERα, PR, retinoic acid (RARα), pregnane X (PXR), and constitutive androstane (CAR) receptors. Therefore, these juvenoids may interfere with key sex steroid hormone receptors, with fenoxycarb having weak antiandrogenic and antiestrogenic properties and pyriproxyfen being antiandrogenic with a weak estrogenic activity. Although these activities appear at the high micromolar concentration range, they pose a potential risk to the human endocrine system.

In this study, we suggest a new mechanism by which juvenoids might interfere with the endocrine system through the AHR, a recognized endocrine target with a wide range of physiological functions. Although this mechanism is plausible given the structural and functional similarities between the JHR and AHR receptors, we did not detect a marked activity of the established juvenoids on human AHR in the reporter assay. However, about a quarter of the JHR agonists newly discovered through our HTS based in *Drosophila* cells modulated AHR activity, suggesting that cross-activities between the insect and the human receptor may be more common than might be expected. While juvenoids offer a promising strategy of controlling insect pests, these results indicate that attention must be paid to prevent undesired endocrine disrupting effects should a new generation of juvenoid insecticides be developed.

## Materials and Methods

### Compounds and chemical library

Fenoxycarb, pyriproxyfen, S-methoprene, S-hydroprene, JH III, B[a]P, TCDD, FICZ and indirubin were purchased from Sigma-Aldrich (Saint Louis, MO, USA); JHB_3_, MF and farnesol were from Echelon Biosciences (Salt Lake City, UT, USA). 10 mM stock solutions in DMSO were prepared, aliquoted and frozen at -20°C. The screening compound library consisting of 50,000 drug-like molecules with molecular weight between 150 and 560 was purchased as individual compounds from Enamine (Kiev, Ukraine) and Chembridge (San Diego, CA, USA). Compounds were dissolved in DMSO to a final concentration of 10 mM, then re-formatted to acoustic dispensing-enabled screening 1536-well plates using automated robotic systems cell∷explorer (PerkinElmer, Waltham, MA, USA) and Vantage (Hamilton, Bonaduz, Switzerland).^61^

### Vectors and stably transfected cell lines for luciferase reporter assays

The firefly luciferase reporter plasmids bearing eight repeats of the wild-type and mutated JH response element from the *Aedes aegypti early trypsin* gene followed by its minimal promoter in the pGL4.17 vector (Promega, Madison, WI, USA) were described earlier.^9^ In this study, they are designated as JHRE [fLuc] and *mut*JHRE [fLuc], respectively (Figure 3 (A)). Additional reporter plasmids, JHRE [nLuc] and *mut*JHRE [nLuc], were prepared by substituting the firefly luciferase sequence with that encoding the NanoLuc® luciferase (Promega, Madison, WI, USA). The NanoLuc® sequence was excised from a previously described vector pNL-9xUAS [nLucP Hygro]^62^ using *Hin*dIII and *Bam*HI, and ligated into JHRE [fLuc] and *mut*JHRE [fLuc] using the same restriction enzymes. EGFP sequence cloned into the pIEx-4 plasmid (Sigma-Aldrich, Saint Louis, MO, USA) served as a negative control for cell line selection.

To establish stable *Drosophila* Kc167 cell lines carrying the JHRE reporter plasmids described above (Figure 3 (A)), we utilized selection for neomycin resistance conferred by the pGL4.17 vector. Briefly, Kc167 cells were seeded in 6-well plates and transfected (3 µg DNA per well) with the JHRE reporter or control (EGFP-pIEx-4) plasmids using the standard FuGENE-HD transfection protocol (Promega, Madison, WI, USA). About 48 h post-transfection, the medium was replaced one containing 900 µg/ml geneticin (G418, Gibco, Thermo Fisher Scientific Inc., Waltham, MA, USA). The medium was changed every three days, keeping the G418 concentration at 900 µg/ml and passaging cells as required. Control cells transfected with EGFP-pIEx-4 died off in about 10 days. The resistant cells containing the reporter plasmids were propagated in the selection medium for additional two weeks. The pool cells were used directly in reporter assays without clonal selection, as isolated clones poorly survived. The clonally selected AZ-AHR cell line was previously described^45^ and kindly provided by Dr. Zdenek Dvorak (Palacky University, Olomouc, Czech Republic). This reporter line is based in human hepatoma HepG2 cells and carries firefly luciferase driven by multiple DREs.

### Cell-based luciferase reporter assays

Technical details of developing reporter assays for compound profiling have been described.^63^ Transcriptional response of JHR or AHR to tested compounds was evaluated using stably transfected *Drosophila* Kc167 reporter cell lines JHRE [fLuc], JHRE [nLuc] and their versions with mutated (*mut*JHRE) as controls. AHR activation was determined in the AZ-AHR cell line.^45^ Kc167 cells were propagated in the Schneider’s *Drosophila* medium (Gibco, 21720-024) supplemented with 10% heat-inactivated fetal bovine serum (Sigma, F4135) and 1x penicillin-streptomycin (Gibco) at 26°C under humidified normal atmosphere. AZ-AHR cells were cultured in the DMEM medium (Gibco, 11880-036) supplemented with 10% heat-inactivated fetal bovine serum, 1 mM non-essential amino acids (Gibco) and 1x penicillin-streptomycin (Gibco) at 37°C under humidified atmosphere with 5% CO_2_.

Cells were harvested, counted and seeded to cell culture-treated, solid white 384- or 1536-well plates (Corning Inc., NY, USA). For Kc167-based reporters, 2,000 or 8,000 cells per well were dispensed to 1536- or 384-well plates in 5 or 20 μl of media, respectively. AZ-AHR cells (5,000 per well) were dispensed to 384-well plates in 20 μl of medium. Cells were dispensed to microtiter plates by Multidrop Combi (Thermo Fisher Scientific Inc., Waltham, MA, USA) or Tempest (Formulatrix LLC, Bedford, MA, USA). Test compounds pre-formatted on 384-well plates as DMSO solutions were transferred to assay plates with contact-free acoustic dispensing technology using ECHO 550 (Beckman Coulter, Inc., Brea, CA, USA) integrated in the fully automated robotic HTS station cell∷explorer (PerkinElmer, Waltham, MA, USA). Plates were vigorously shaken for 60 s and then incubated in the humidified cell incubator overnight. The reporter activity was determined by dispensing 2.5 μl or 12 μl of luciferase reagents per well in 1536- or 384-well plates, respectively. Bright-Glo™ Luciferase Assay System (Promega, Madison, WI, USA) was used for the firefly-based reporters and Nano-Glo® Luciferase Assay System (Promega, Madison, WI, USA) for NanoLuc® luciferase-based reporter assays. After shaking for 2 min, plates were centrifuged and luminescence was recorded on the Envision plate reader (PerkinElmer, Waltham, MA, USA).

Data were collected, processed, and normalized using the LIMS system ScreenX.^64^ Luciferase activities were normalized on the scale from 0% to 100%, where 0% corresponds to the activity of cells treated with DMSO only and 100% corresponds to the reporter activity obtained with the saturating concentration of the reference agonist.

### Cell viability assay

Cell viability was monitored along with the reporter assays using the same plate type and format, equal number of cells per well, and the same incubation protocol with the tested compounds. Upon incubation, intracellular ATP levels were quantified using the CellTiter-Glo 2.0 luminescence assay (Promega, Madison, WI, USA). Luciferase activity was recorded on the Envision plate reader (PerkinElmer, Waltham, MA, USA) and normalized on a scale from 0 to 100% (0%, activity in the growth medium without cells; 100%, activity in cells treated with DMSO only).

### High-throughput screening of chemical library

DMSO solutions (10 mM) of 50,000 compounds preformatted on 1536-well plates were transferred to empty white, cell-culture treated, 1536-well plates (Corning, NY, USA) using the acoustic dispensing technology as described above. Upon transfer, each plate was protected with foil against evaporation. Stably transfected JHRE [fLuc] reporter cells were grown as described above, harvested, counted and seeded to the plates containing the pre-dispensed chemical library. To each well, 2,000 cells were dispensed in 5 μl of medium using Multidrop Combi (Thermo Fisher Scientific Inc., Waltham, MA, USA). The final concentration of the screened compounds was 10 μM. Each plate was shaken for 60 s using a Variomag shaker (Thermo Fisher Scientific Inc., Waltham, MA, USA) and then incubated at 37°C under 5% CO_2_. The firefly luciferase activity was determined 24 h later with the Bright-Glo™ Luciferase Assay System (Promega, Madison, WI, USA) on the Envision plate reader (PerkinElmer, Waltham, MA, USA) following the manufacturer’s protocol.

### JH binding assays

The DNA sequence encoding *Drosophila melanogaster* GCE (NCBI Accession <u>NP_001188593.1</u>) cloned in the *pK-Myc-C2* vector was a template for site-directed mutagenesis by PCR with overlapping primers to generate the individual Y540F, T542Y, Y604F, and S618A amino acid substitutions. The wild-type and mutated GCE variants were expressed *in vitro* using the TnT Quick T7 Coupled translation rabbit reticulocyte lysate system (Promega, Madison, WI, USA). The reaction products were examined in the dextran-coated charcoal binding assay^4,9^ with own synthesized [^3^H]JH III (25.7 Ci/mmol).^62^ The reticulocyte lysate without DNA input was the background control.

## Data analysis

Data from plate-based cellular assays were uploaded to the proprietary LIMS system ScreenX.^64^ There, data were further processed by internal algorithm and associated with corresponding chemical structures. Data from the primary HTS were normalized using the B-score algorithm^46^ to eliminate artifacts arising from the location of the sample on the microtiter plate, such as the edge effect or gradients, as well as any systematic errors introduced by the automated devices during the HTS process, including row and column effects caused by bulk-dispensers. Compounds showing B-score activities > 5 were marked as hits and were further validated in dose-dependent studies. Compound structures were drawn in Marvin chemical editor (version 23.11, Chemaxon, Budapest, Hungary). EC_50_ values were obtained using nonlinear regression analysis (Y=Bottom + (Top-Bottom)/(1+10((LogEC50-X) *HillSlope)) by GraphPad Prism (Version 10, Boston, MA., USA). The GraphPad Prism software was also used for graph preparation. The chemical analysis, calculation of cLogP, was performed by TIBCO Spotfire® Analyst (version 12.0.3.77, Palo Alto, CA, USA) expanded with Lead Discovery Premium cheminformatics platform by Perkin Elmer (Waltham, MA, USA). Calculations of physico-chemical properties such as polar surface area and van der Waals volume were carried out with Calculators & Predictors (version 23.11, Chemaxon, Budapest, Hungary).

### Molecular modeling

Ligand docking and MD simulations were performed as previously described^32^ using GROMACS,^65^ AMBER99SB-ILDN force field^66^ supplemented with generalized AMBER force field parameters^67^ and the TIP3P explicit water model. Modified Berendsen thermostat^68^ was used to maintain constant temperature 300 K while Parrinello-Rahman coupling^69^ was employed to keep pressure constant at 1 bar using an octagonal box. System charge was neutralized by chloride or sodium ions. Simulations were performed for 1 μs or until stable root mean square deviation (RMSD) was maintained for at least 200 ns. In order to select representative, high-affinity frames, MD trajectories were sampled every 1 ns and their energy was computed using the GBSA approximation Still’s model^70^ for computation of the effective Born radii as previously described.^32^ CAVER software was used for static cavity evaluation and visualization^71^ while dynamic cavity volume averaging was done using trj_cavity.^72^

### CRediT authorship contribution statement

**David Sedlak**: Conceptualization, Methodology, Formal analysis, Investigation, Data curation, Writing - Original draft, Writing - Review & editing, Visualization, Supervision, Project administration, Funding acquisition. **Roman Tuma**: Conceptualization, Methodology, Formal analysis, Investigation, Data curation, Writing - Original draft, Writing - Review & editing, Visualization. **Jayaprakash Narayana Kolla**: Formal analysis, Investigation. **Raveendra Babu Mokhamatam**: Methodology, Formal analysis, Investigation, Data curation. **Liliia Bahrova**: Formal analysis, Investigation. **Michaela Lisova**: Investigation. **Lenka Bittova**: Formal analysis, Investigation. **Marek Jindra**: Conceptualization, Methodology, Investigation, Writing - Original draft, Writing - Review & editing, Supervision, Project administration, Funding acquisition.

## Abbreviations

AHR: aryl hydrocarbon receptor
ARNT: aryl hydrocarbon receptor nuclear translocator
B[a]P: benzo[a]pyrene
bHLH-PAS: basic helix-loop-helix PER-ARNT-SIM
cryo-EM: cryogenic electron microscopy
DRE: dioxin response element
EDC: endocrine disruptive compound
ER: estrogen receptor
FICZ: 6-formylindolo[3,2-b]carbazole
GBSA: generalized Born surface area
GCE: germ cell-expressed
HSP90: heat-shock protein 90
HTS: high-throughput screening
JH III: juvenile hormone III
JHB_3_: JH III bisepoxide
JHR: juvenile hormone receptor
JHRE: juvenile hormone response element
mAHR: murine AHR
MD: molecular dynamics
MET: methoprene-tolerant
MF: methyl farnesoate
NCoA: nuclear receptor coactivator
NR: nuclear receptor
PAH: polycyclic aromatic hydrocarbon
RMSD: root mean square deviation
TAI: taiman
TCDD: 2,3,7,8-tetrachlorodibenzo-*p*-dioxin
XAP2: hepatitis B virus protein X-associated protein.

## Acknowledgments

We highly appreciate the help of Olga Martinkova in cell culturing and preparation of stable cell lines, Martin Popr and Tomas Langhammer for help with compound management and lab automation, respectively, and Tomas Muller for cheminformatics support (ScreenX). We are grateful to Zdenek Dvorak for his generous gift of the clonal AZ-AHR cell line. This research was supported by grant project 20-05151X from the Czech Science Foundation to MJ and DS. Project LM2018130 from the Czech Ministry of Education, Youth, and Sports (MEYS) and the National Program for Sustainability (RVO: 68378050-KAV-NPUI) from the Czech Academy of Sciences funded the National Infrastructure CZ-OPENSCREEN, led by Dr. Petr Bartunek. RT and computational resources were supported by ERDF project CZ.02.1.01/0.0/0.0/15_003/0000441.

## Notes

### Competing Interest Statement

David Sedlak and Marek Jindra have co-founded a startup company Preagon Biotech.

### Summary of Updates

The secondary affiliation of D. Sedlak and M. Jindra to Preagon Biotech has been placed in the footnote along with current e-mail address of D. Sedlak. Changes are limited to the title pages of the main text and the SI.

## References

1. Jindra, M., Palli, S.R. and Riddiford, L.M. (2013) The juvenile hormone signaling pathway in insect development. Annu. Rev. Entomol. 58, 181–204.

2. Roy, S., Saha, T.T., Zou, Z. and Raikhel, A.S. (2018) Regulatory pathways controlling female insect reproduction. Annu. Rev. Entomol. 63, 489–511.

3. Ashok, M., Turner, C. and Wilson, T.G. (1998) Insect juvenile hormone resistance gene homology with the bHLH-PAS family of transcriptional regulators. Proc. Natl. Acad. Sci. USA 95, 2761–2766.

4. Charles, J.P., Iwema, T., Epa, V.C., Takaki, K., Rynes, J. and Jindra, M. (2011) Ligand-binding properties of a juvenile hormone receptor, Methoprene-tolerant. Proc. Natl. Acad. Sci. USA 108, 21128–21133.

5. Jindra, M., Tumova, S., Milacek, M., Bittova, L. (2021) A decade with the juvenile hormone receptor. Adv. Insect Physiol. 60, 37–85.

6. Wilson, T.G. and Fabian, J. (1986) A *Drosophila melanogaster* mutant resistant to a chemical analog of juvenile hormone. Dev. Biol. 118, 190–201.

7. Baumann, A., Fujiwara, Y. and Wilson, T.G. (2010) Evolutionary divergence of the paralogs Methoprene tolerant (Met) and germ cell expressed (gce) within the genus *Drosophila*. J. Insect Physiol. 56, 1445–1455.

8. Tumova, S., Dolezel, D. and Jindra, M. (2023) Conserved and unique roles of bHLH-PAS transcription factors in insects - From clock to hormone reception. J. Mol. Biol. 168332. doi: 10.1016/j.jmb.2023.168332

9. Jindra, M., Uhlirova, M., Charles, J.P., Smykal, V. and Hill, R.J. (2015) Genetic evidence for function of the bHLH-PAS protein Gce/Met as a juvenile hormone receptor. PLoS Genet. 11, e1005394.

10. Abdou, M.A., He, Q., Wen, D., Zyaan, O., Wang, J., Xu, J., Baumann, A.A., Joseph, J., Wilson, T.G., Li, S. et al. (2011) *Drosophila* Met and Gce are partially redundant in transducing juvenile hormone action. Insect Biochem. Mol. Biol. 41, 938–945.

11. Baumann, A.A., Texada, M.J., Chen, H.M., Etheredge, J.N., Miller, D.L., Picard, S., Warner, R., Truman, J.W. and Riddiford, L.M. (2017) Genetic tools to study juvenile hormone action in *Drosophila*. Sci. Rep. 7, 2132.

12. Klonaros, D., Dresch, J.M. and Drewell, R.A. (2023) Transcriptome profile in *Drosophila* Kc and S2 embryonic cell lines. G3 13, jkad054.

13. Li, M., Mead, E.A. and Zhu, J. (2011) Heterodimer of two bHLH-PAS proteins mediates juvenile hormone-induced gene expression. Proc. Natl. Acad. Sci. USA 108, 638–643.

14. Li, M., Liu, P., Wiley, J.D., Ojani, R., Bevan, D.R., Li, J. and Zhu, J. (2014) A steroid receptor coactivator acts as the DNA-binding partner of the methoprene-tolerant protein in regulating juvenile hormone response genes. Mol. Cell. Endocrinol. 394, 47–58.

15. He, Q., Wen, D., Jia, Q., Cui, C., Wang, J., Palli, S.R. and Li, S. (2014) Heat shock protein 83 (Hsp83) facilitates methoprene-tolerant (Met) nuclear import to modulate juvenile hormone signaling. J. Biol. Chem. 289, 27874–27885.

16. Jindra, M., McKinstry, W.J., Nebl, T., Bittova, L., Ren, B., Shaw, J., Phan, T., Lu, L., Low, J.K.K., Mackay, J.P. et al. (2021) Purification of an insect juvenile hormone receptor complex enables insights into its post-translational phosphorylation. J. Biol. Chem. 297, 101387.

17. Tumova, S. and Jindra, M. (2023) Ligand-dependent protein interactions of the juvenile hormone receptor captured in real time. FEBS J. 290, 2881–2894.

18. Kayukawa, T., Minakuchi, C., Namiki, T., Togawa, T., Yoshiyama, M., Kamimura, M., Mita, K., Imanishi, S., Kiuchi, M., Ishikawa, Y. et al. (2012) Transcriptional regulation of juvenile hormone mediated induction of Kruppel homolog 1, a repressor of insect metamorphosis. Proc. Natl. Acad. Sci. USA 109, 11729–11734.

19. Miyakawa, H. and Iguchi, T. (2017) Comparative luciferase assay for establishing reliable in vitro screening system of juvenile hormone agonists. J. Appl. Toxicol. 37, 1082–1090.

20. Kewley, R.J., Whitelaw, M.L. and Chapman-Smith, A. (2004) The mammalian basic helix-loop-helix/PAS family of transcriptional regulators. Int. J. Biochem. Cell Biol. 36, 189–204.

21. Beischlag, T.V., Luis Morales, J., Hollingshead, B.D. and Perdew, G.H. (2008) The aryl hydrocarbon receptor complex and the control of gene expression. Crit. Rev. Eukaryot. Gene Expr. 18, 207–250.

22. Denison, M.S. and Faber, S.C. (2017) And now for something completely different: Diversity in ligand-dependent activation of Ah receptor responses. Curr. Opin. Toxicol. 2, 124–131.

23. Murray, I.A., Morales, J.L., Flaveny, C.A., Dinatale, B.C., Chiaro, C., Gowdahalli, K., Amin, S. and Perdew, G.H. (2010) Evidence for ligand-mediated selective modulation of aryl hydrocarbon receptor activity. Mol. Pharmacol. 77, 247–254.

24. Sladekova, L., Mani, S. and Dvorak, Z. (2023) Ligands and agonists of the aryl hydrocarbon receptor AhR: Facts and myths. Biochem. Pharmacol. 213, 115626.

25. Bittova, L., Jedlicka, P., Dracinsky, M., Kirubakaran, P., Vondrasek, J., Hanus, R. and Jindra, M. (2019) Exquisite ligand stereoselectivity of a *Drosophila* juvenile hormone receptor contrasts with its broad agonist repertoire. J. Biol. Chem. 294, 410–423.

26. Jindra, M. and Bittova, L. (2020) The juvenile hormone receptor as a target of juvenoid “insect growth regulators”. Arch. Insect Biochem. Physiol. 103, e21615.

27. Gruszczyk, J., Grandvuillemin, L., Lai-Kee-Him, J., Paloni, M., Savva, C.G., Germain, P., Grimaldi, M., Boulahtouf, A., Kwong, H.S., Bous, J. et al. (2022) Cryo-EM structure of the agonist-bound Hsp90-XAP2-AHR cytosolic complex. Nat. Commun. 13, 7010.

28. Denison, M.S. (2010). National Center for Biotechnology Information. PubChem Bioassay Record for AID 602173, Source: The Scripps Research Institute Molecular Screening Center. https://pubchem.ncbi.nlm.nih.gov/bioassay/602173.

29. Kayukawa, T., Furuta, K., Nagamine, K., Shinoda, T., Yonesu, K. and Okabe, T. (2020) Identification of a juvenile-hormone signaling inhibitor via high-throughput screening of a chemical library. Sci. Rep. 10, 18413.

30. Kayukawa, T., Furuta, K., Yonesu, K. and Okabe, T. (2021) Identification of novel juvenile-hormone signaling activators via high-throughput screening with a chemical library. J. Pestic. Sci. 46, 53–59.

31. Yokoi, T., Nabe, T., Horoiwa, S., Hayashi, K., Ito-Harashima, S., Yagi, T., Nakagawa, Y. and Miyagawa, H. (2021) Virtual screening identifies a novel piperazine-based insect juvenile hormone agonist. J. Pestic. Sci. 46, 68–74.

32. Tumova, S., Milacek, M., Snajdr, I., Muthu, M., Tuma, R., Reha, D., Jedlicka, P., Bittova, L., Novotna, A., Majer, P. et al. (2022) Unique peptidic agonists of a juvenile hormone receptor with species-specific effects on insect development and reproduction. Proc. Natl. Acad. Sci. USA 119, e2215541119.

33. Devillers, J. (2020) Fate and ecotoxicological effects of pyriproxyfen in aquatic ecosystems. Environ. Sci. Pollut. Res. Int. 27, 16052–16068.

34. Cuvillier-Hot, V. and Lenoir, A. (2020) Invertebrates facing environmental contamination by endocrine disruptors: Novel evidences and recent insights. Mol. Cell. Endocrinol. 504, 110712.

35. Toyota, K., Watanabe, H., Hirano, M., Abe, R., Miyakawa, H., Song, Y., Sato, T., Miyagawa, S., Tollefsen, K.E., Yamamoto, H. et al. (2022) Juvenile hormone synthesis and signaling disruption triggering male offspring induction and population decline in cladocerans (water flea): Review and adverse outcome pathway development. Aquat. Toxicol. 243, 106058.

36. Crane, M., Dungey, S., Lillicrap, A., Thompson, H., Weltje, L., Wheeler, J.R. and Lagadic, L. (2022) Commentary: Assessing the endocrine disrupting effects of chemicals on invertebrates in the European Union. Environ. Sci. Eur. 34, 36.

37. Balaguer, P., Delfosse, V. and Bourguet, W. (2019) Mechanisms of endocrine disruption through nuclear receptors and related pathways. Curr. Opin. Endocr. Metab. Res. 7, 1–8.

38. Kojima, H., Takeuchi, S. and Nagai, T. (2010) Endocrine-disrupting potential of pesticides via nuclear receptors and aryl hydrocarbon receptor. J. Health Sci. 56, 374–386.

39. Seok, S.H., Lee, W., Jiang, L., Molugu, K., Zheng, A., Li, Y., Park, S., Bradfield, C.A. and Xing, Y. (2017) Structural hierarchy controlling dimerization and target DNA recognition in the AHR transcriptional complex. Proc. Natl. Acad. Sci. USA 114, 5431–5436.

40. Schulte, K.W., Green, E., Wilz, A., Platten, M. and Daumke, O. (2017) Structural basis for aryl hydrocarbon receptor-mediated gene activation. Structure 25, 1025–1033.e3.

41. Jumper, J., Evans, R., Pritzel, A., Green, T., Figurnov, M., Ronneberger, O., Tunyasuvunakool, K., Bates, R., Zidek, A., Potapenko, A. et al. (2021) Highly accurate protein structure prediction with AlphaFold. Nature 596, 583–589.

42. Pandini, A., Soshilov, A.A., Song, Y., Zhao, J., Bonati, L. and Denison, M.S. (2009) Detection of the TCDD binding-fingerprint within the Ah receptor ligand binding domain by structurally driven mutagenesis and functional analysis. Biochemistry 48, 5972–5983.

43. Motto, I., Bordogna, A., Soshilov, A.A., Denison, M.S. and Bonati, L. (2011) New aryl hydrocarbon receptor homology model targeted to improve docking reliability. J. Chem. Inf. Model. 51, 2868–2881.

44. Wen, Z., Zhang, Y., Zhang, B., Hang, Y., Xu, L., Chen, Y., Xie, Q., Zhao, Q., Zhang, L., Li, G. et al. (2023) Cryo-EM structure of the cytosolic AhR complex. Structure 31, 295–308.e4.

45. Novotna, A., Pavek, P. and Dvorak, Z. (2011) Novel stably transfected gene reporter human hepatoma cell line for assessment of aryl hydrocarbon receptor transcriptional activity: construction and characterization. Environ. Sci. Technol. 45, 10133–10139.

46. Brideau, C., Gunter, B., Pikounis, B. and Liaw, A. (2003) Improved statistical methods for hit selection in high-throughput screening. J. Biomol. Screen. 8, 634–647.

47. Han, D., Nagy, S.R. and Denison, M.S. (2004) Comparison of recombinant cell bioassays for the detection of Ah receptor agonists. Biofactors 20, 11–22.

48. Pener, M.P., Dhadialla, T.S. (2012) An overview of insect growth disruptors; applied aspects. Adv. Insect Physiol. 43, 1–162.

49. Parthasarathy, R., Farkas, R. and Palli, S.R. (2012) Recent progress in juvenile hormone analogs (JHA) research. Adv. Insect Physiol. 43, 353–436.

50. Heath-Pagliuso, S., Rogers, W.J., Tullis, K., Seidel, S.D., Cenijn, P.H., Brouwer, A. and Denison, M.S. (1998) Activation of the Ah receptor by tryptophan and tryptophan metabolites. Biochemistry 37, 11508–11515.

51. Zelante, T., Iannitti, R.G., Cunha, C., De Luca, A., Giovannini, G., Pieraccini, G., Zecchi, R., D’Angelo, C., Massi-Benedetti, C., Fallarino, F., et al. (2013) Tryptophan catabolites from microbiota engage aryl hydrocarbon receptor and balance mucosal reactivity via interleukin-22. Immunity 39, 372–385.

52. Schroeder, J.C., DiNatale, B.C., Murray, I.A., Flaveny, C.A., Liu, Q., Laurenzana, E.M., Lin, J.M., Strom, S.C., Omiecinski, C.J., Amin, S. et al. (2010) The uremic toxin 3-indoxyl sulfate is a potent endogenous agonist for the human aryl hydrocarbon receptor. Biochemistry 49, 393–400.

53. Bjeldanes, L.F., Kim, J.Y., Grose, K.R., Bartholomew, J.C. and Bradfield, C. A. (1991) Aromatic hydrocarbon responsiveness-receptor agonists generated from indole-3-carbinol in vitro and in vivo: comparisons with 2,3,7,8-tetrachlorodibenzo-p-dioxin. Proc. Natl. Acad. Sci. USA 88, 9543–9547.

54. Vyhlídalová, B., Krasulová, K., Pečinková, P., Poulíková, K., Vrzal, R., Andrysík, Z., Chandran, A., Mani, S. and Dvorak, Z. (2020) Antimigraine drug avitriptan is a ligand and agonist of human aryl hydrocarbon receptor that induces CYP1A1 in hepatic and intestinal cells. Int. J. Mol. Sci. 21, 2799.

55. Casals-Casas, C. and Desvergne, B. (2011) Endocrine disruptors: from endocrine to metabolic disruption. Annu. Rev. Physiol. 73, 135–162.

56. Ohtake, F., Takeyama, K., Matsumoto, T., Kitagawa, H., Yamamoto, Y., Nohara, K., Tohyama, C., Krust, A., Mimura, J., Chambon, P. et al. (2003) Modulation of oestrogen receptor signalling by association with the activated dioxin receptor. Nature 423, 545–550.

57. Delfosse, V., Maire, A.L., Balaguer, P. and Bourguet, W. (2015) A structural perspective on nuclear receptors as targets of environmental compounds. Acta Pharmacol. Sin. 36, 88–101.

58. La Merrill, M.A., Vandenberg, L.N., Smith, M.T., Goodson, W., Browne, P., Patisaul, H.B., Guyton, K.Z., Kortenkamp, A., Cogliano, V.J., Woodruff, T.J., et al. (2020) Consensus on the key characteristics of endocrine-disrupting chemicals as a basis for hazard identification. Nat. Rev. Endocrinol. 16, 45–57.

59. Richard, A.M., Huang, R., Waidyanatha, S., Shinn, P., Collins, B.J., Thillainadarajah, I., Grulke, C.M., Williams, A.J., Lougee, R.R., Judson, R.S. et al. (2021) The Tox21 10K compound library: Collaborative chemistry advancing toxicology. Chem. Res. Toxicol. 34, 189–216.

60. Ngan, D.K., Xia, M., Simeonov, A. and Huang, R. (2023) In vitro profiling of pesticides within the Tox21 10K compound library for bioactivity and potential toxicity. Toxicol. Appl. Pharmacol. 473, 116600.

61. Popr, M., Sedlak, D. and Bartunek, P. (2017) Management - maintenance of chemical compound libraries for use in high throughput screening. Chem. Listy 111, 727–776.

62. Milacek, M., Bittova, L., Tumova, S., Luksan, O., Hanus, R., Kyjakova, P., Machara, A., Marek, A. and Jindra, M. (2021) Binding of de novo synthesized radiolabeled juvenile hormone (JH III) by JH receptors from the Cuban subterranean termite *Prorhinotermes simplex* and the German cockroach *Blattella germanica*. Insect Biochem. Mol. Biol. 139, 103671.

63. Sedlak, D., Paguio, A. and Bartunek, P. (2011) Two panels of steroid receptor luciferase reporter cell lines for compound profiling. Comb. Chem. High Throughput Screen. 14, 248–266.

64. Muller, T., Sedlak, D. and Bartunek, P. (2017) Laboratory information systems to high-throughput screening. Chem. Listy 111, 766–771.

65. Van Der Spoel, D., Lindahl, E., Hess, B., Groenhof, G., Mark, A.E. and Berendsen, H.J. (2005) GROMACS: fast, flexible, and free. J. Comput. Chem. 26, 1701–1718.

66. Hornak, V., Abel, R., Okur, A., Strockbine, B., Roitberg, A. and Simmerling, C. (2006) Comparison of multiple amber force fields and development of improved protein backbone parameters. Proteins 65, 712–725.

67. Wang, J., Wolf, R.M., Caldwell, J.W., Kollman, P.A. and Case, D.A. (2004) Development and testing of a general amber force field. J. Comput. Chem. 25, 1157–1174.

68. Berendsen, H.J.C., Postma, J.P.M., van Gunsteren, W.F. A. D. and Haak, J.R. (1984) Molecular dynamics with coupling to an external bath *J*. Chem. Phys. 81, 3684–3690.

69. Parrinello, M. and Rahman, A. (1981) Polymorphic transitions in single crystals: A new molecular dynamics method. J. Appl. Phys. 52, 7182–7190.

70. Still, W.C., Tempczyk, A., Hawley, R.C. and Hendrickson, T. (1990) Semianalytical treatment of solvation for molecular mechanics and dynamics. J. Am. Chem. Soc. 112, 6127–6129.

71. Jurcik, A., Bednar, D., Byska, J., Marques, S.M., Furmanova, K., Daniel, L., Kokkonen, P., Brezovsky, J., Strnad, O., Stourac, J. et al. (2018) CAVER Analyst 2.0: analysis and visualization of channels and tunnels in protein structures and molecular dynamics trajectories. Bioinformatics 34, 3586–3588.

72. Paramo, T., East, A., Garzón, D., Ulmschneider, M.B. and Bond, P.J. (2014) Efficient characterization of protein cavities within molecular simulation trajectories: trj_cavity. J. Chem. Theory Comput. 10, 2151–2164.

