## Supplementary Material for "Unique and Common Agonists Activate the Insect Juvenile Hormone Receptor and the Human AHR"

### for

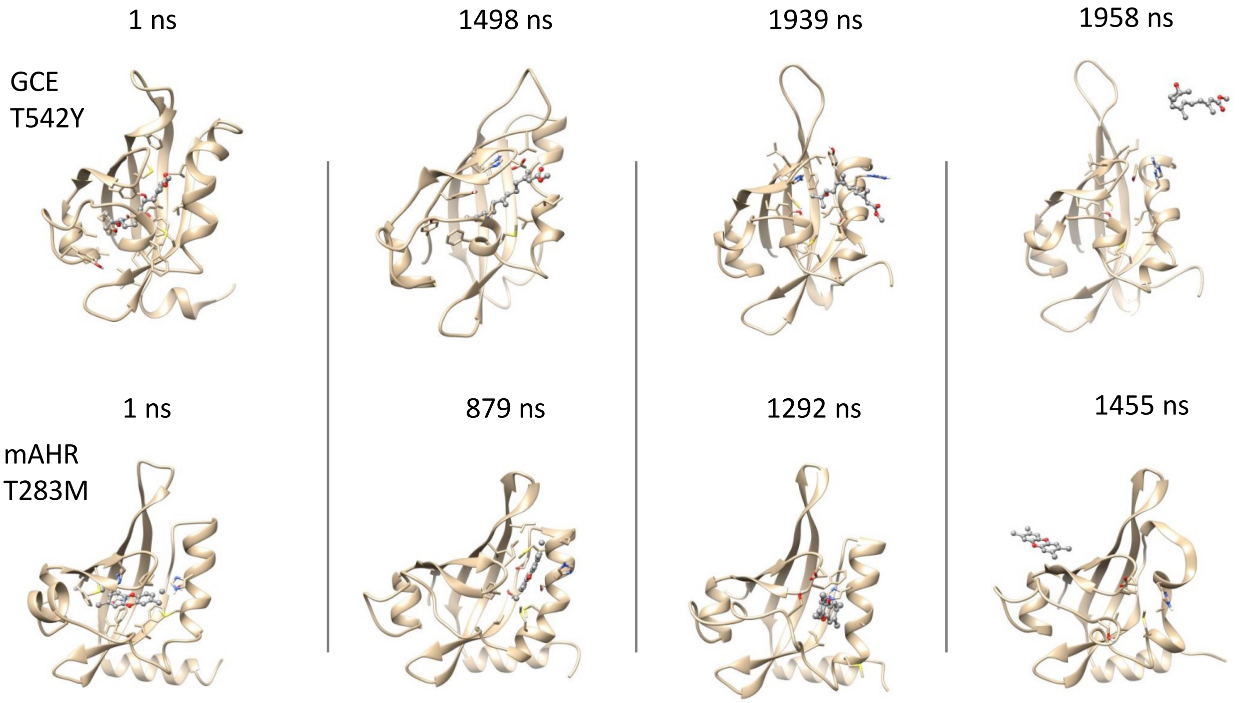


**Figure S1**. Snapshots of MD simulation trajectories sampling ligand exit from mutant receptor PAS-B domains. Panels from the left to the right illustrate the exit of the respective ligand (ball & stick representation) from the binding pocket (ribbon representation of backbone with contact residues shown in stick representation). Due to stochastic nature of the process, different trajectories exhibit exit positions at different times and in many cases no exit occurred within the 3-μs simulation. Top: GCE T542Y mutant with JH III positioned into the binding pocked according to the equilibrated model of the wild-type PAS-B (based on cryo-EM structure PDB 8H77[1]). Bottom: murine AHR T289M mutant with TCDD placed into the PAS-B domain.

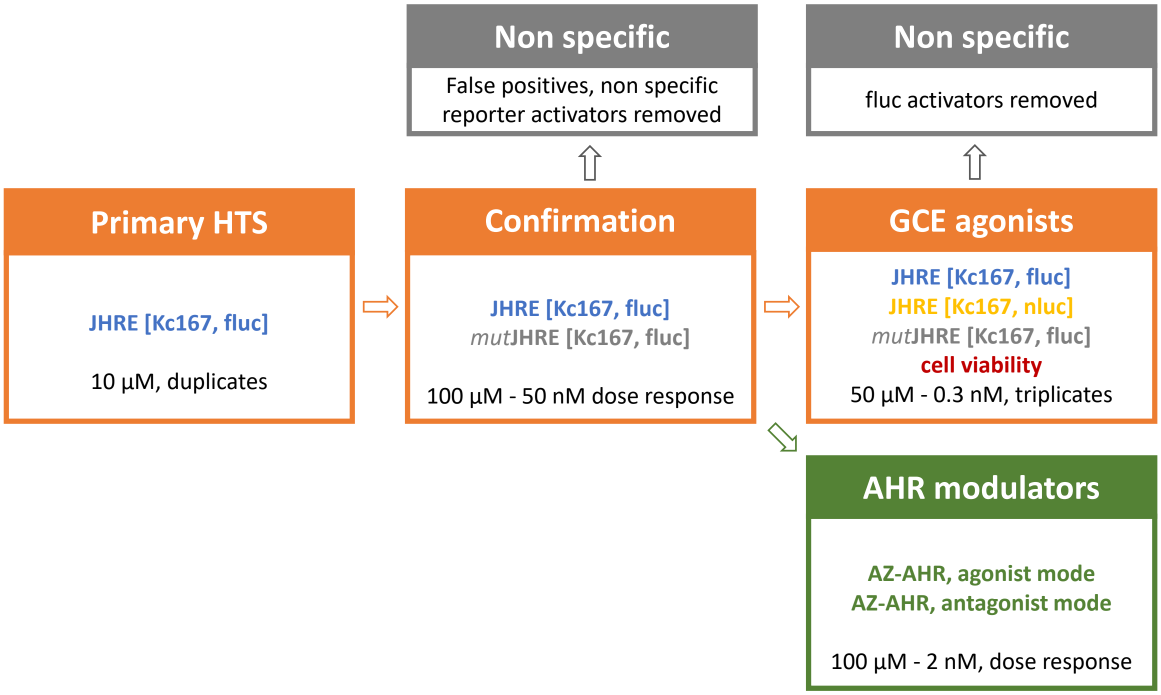


**Figure S2**. Flow chart presenting the HTS process for GCE and AHR unique and common modulators. The screening for GCE agonists was performed with 50,000 compounds at a concentration of 10 µM, with each compound being tested in duplicate. False positives and nonspecific reporter activators were eliminated in the following confirmation step where each hit was retested in a dose-dependent manner using the same reporter assay and the reporter assay containing mutated JHRE (*mut*JHRE). The active compounds were fully characterized with the JHR reporter assays, employing two different luciferases, firefly and NanoLuc®, to determine $EC_{50}$ and to eliminate firefly luciferase activators. JHR agonists were further evaluated in dose-response experiments for AHR agonistic and antagonistic activities. Similar to the reporter assays, each active compound was assessed for its potential cytotoxic effect in a cell viability experiment.

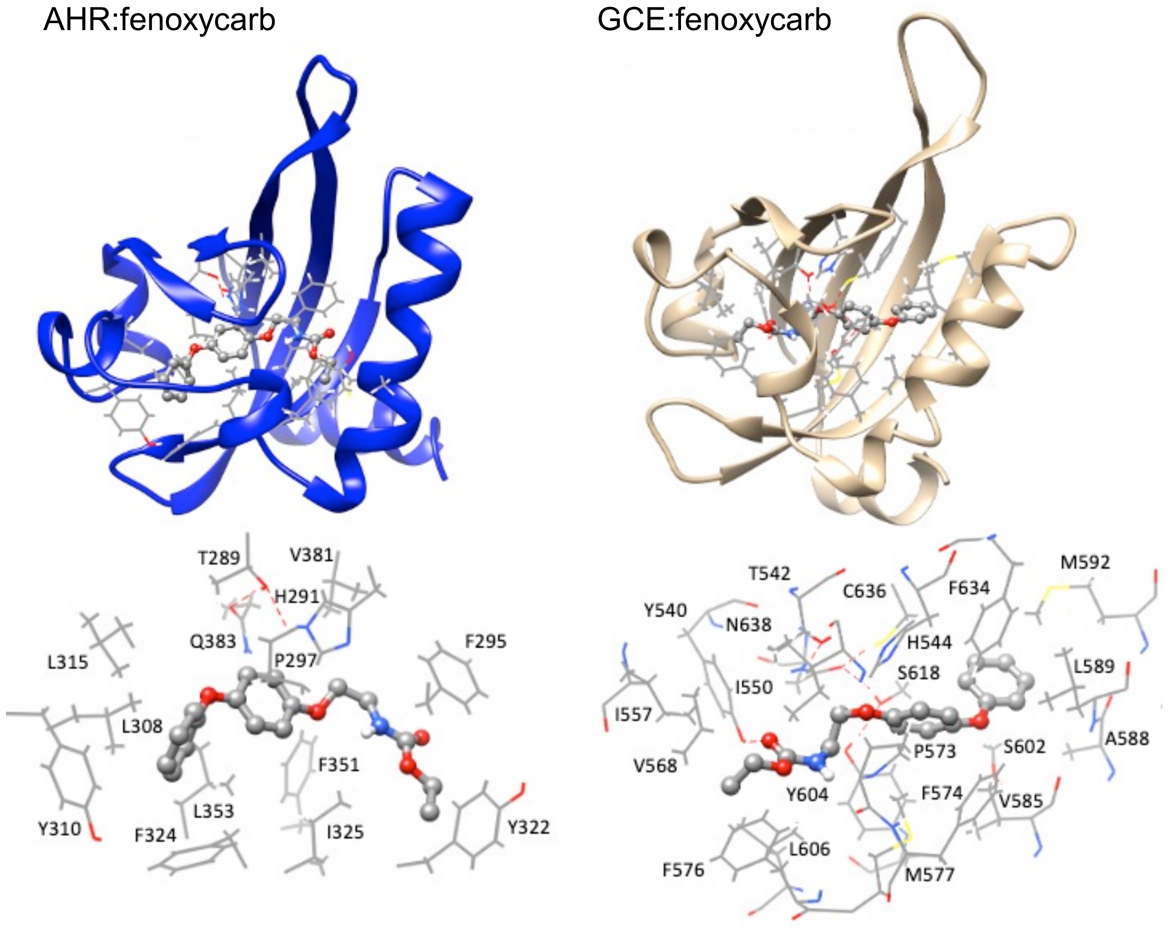


**Figure S3**. Predicted interactions of AHR and GCE PAS-B domains with the GCE ligand fenoxycarb. Top: the AHR and GCE PAS-B models with bound fenoxycarb are shown as blue and wheat ribbons, respectively. The bottom panels depict details of interacting residues (wire) within the MD frames exhibiting the strongest interactions; fenoxycarb is shown in ball & stick representation. Hydrogen bonds are delineated in red dashed lines.

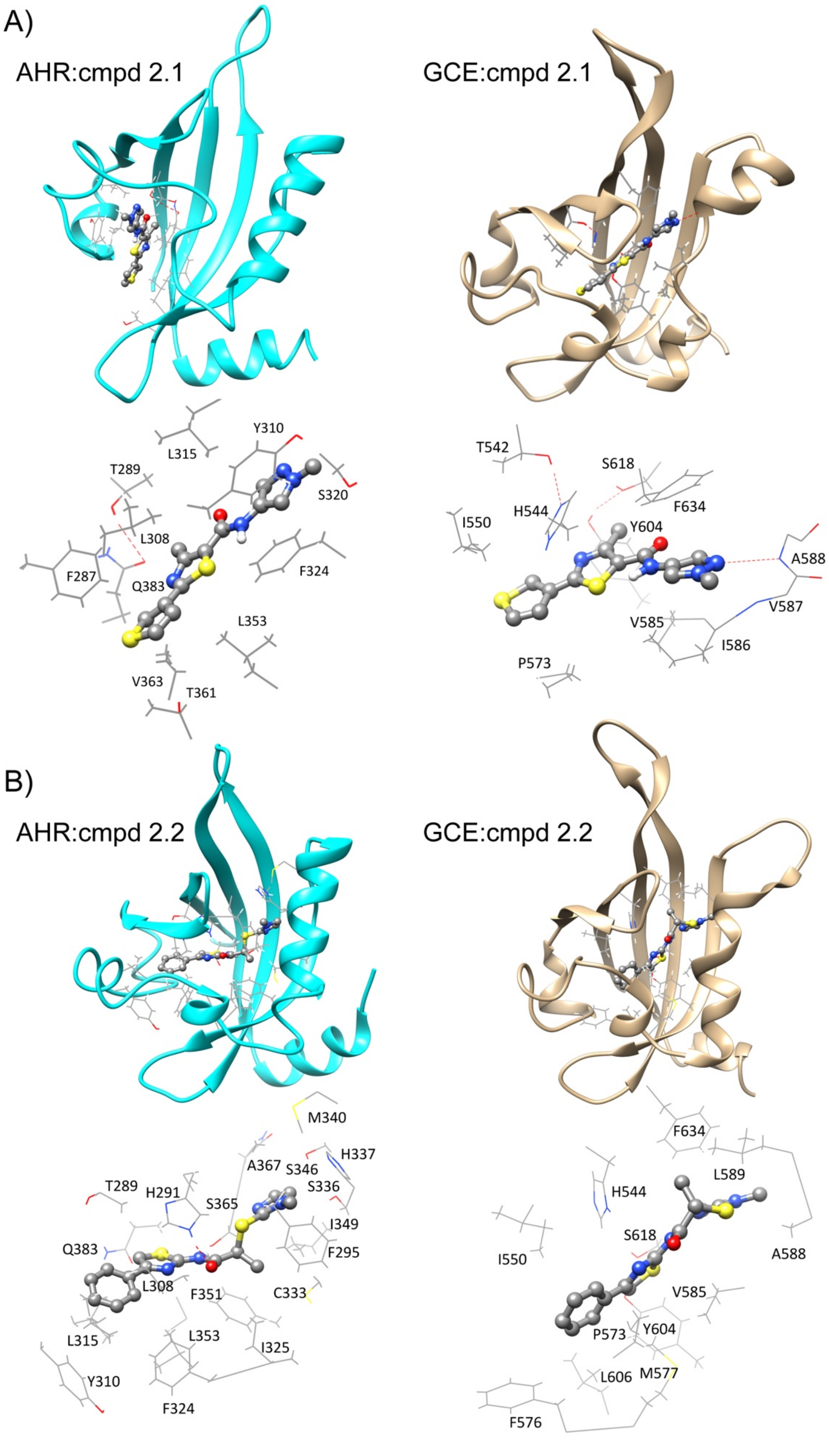
A)
AHR:cmpd 2.1
GCE:cmpd 2.1
L315
Y310
T289
S320
L308
F287
Q383
F324
L353
V363
T361
T542
S618
H544
F634
I550
Y604
A588
V587
V585
I586
P573
B)
AHR:cmpd 2.2
GCE:cmpd 2.2
M340
H337
A367
S346
T289
H291
S336
S365
I349
Q383
F295
L308
F351
C333
L315
L353
I325
Y310
F324
F634
L589
H544
S618
I550
A588
V585
P573
Y604
L606
M577
F576

**Figure S4**. Predicted interactions of AHR and GCE with their common novel agonists cmpds 2.1 (A) and 2.2 (B). GCE and AHR PAS-B models with the bound agonists are shown as cyan and wheat ribbons, respectively. The bottom panels depict details of interacting residues (wire) within the MD frames exhibiting the strongest interactions; cmpds 2.1 and 2.2 are shown in ball & stick representation. Hydrogen bonds are delineated in red dashed lines.

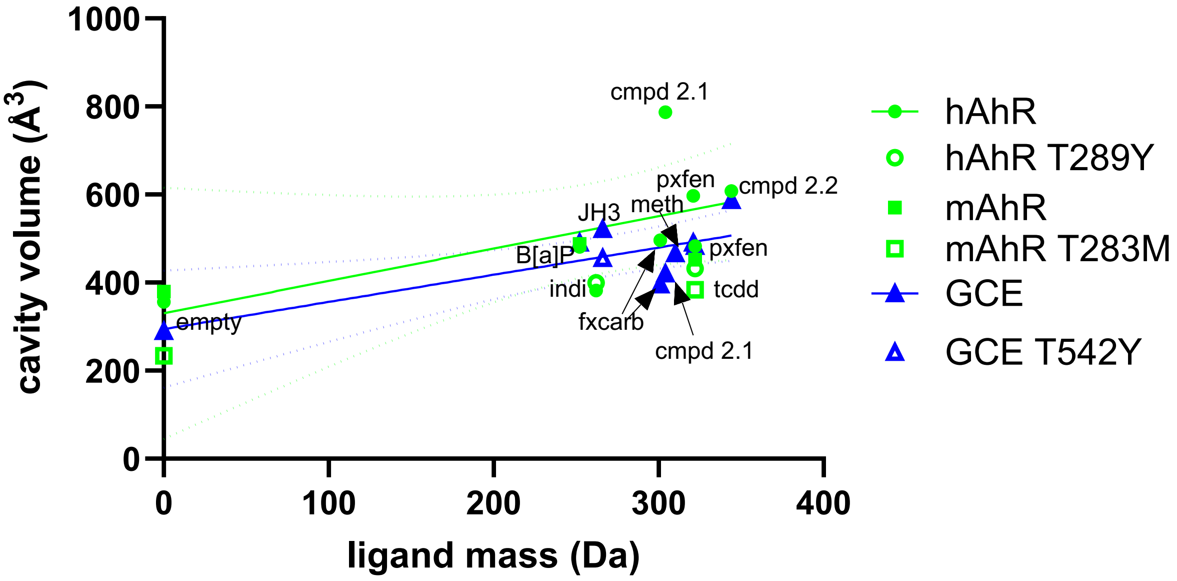


**Figure S5**. Response of the average cavity size to the ligand mass. The averages were computed over the stationary part of 1.5-μs MD trajectory using trj_cavity tool.[2] The models of human AHR (hAHR) from PDB 7ZUB[3] (filled green circles) and GCE PAS-B (blue triangles) were used as starting structures for simulations. Murine AHR (mAHR) model of empty PAS-B from PDB 8H77[1] (green square) and several mutants were used as controls (open symbols). Linear regression (solid lines with 95% confidence intervals shown in dotted lines) was used to test any significant trend for cavity size with respect to the ligand mass. Selected ligands: B[a]P; indi, indirubin; JH3, JH III; fxcarb, fenoxycarb; pxfen, pyriproxyfen; meth, S-methoprene. Empty structures were assigned ligand mass 0. Note that the trajectory averaged cavity sizes do not necessarily match those reported for the representative frames because of large fluctuations in the cavity size and different algorithms.

**Table S1** 93 agonists discovered in the HTS and their activities in the JHR and AHR reporter assays

| **Compound** | JHR $EC_{50}$ [μM] | Efficacy [%][a] | AHR $EC_{50}$ [μM] | Efficacy [%][b] | Mode[c] | MW | cLogP[d] |
|---|---|---|---|---|---|---|---|
| fenoxycarb | 0.05 | 100 | n.a. | n.a. | n.a. | 301 | 3.6 |
| pyriproxyfen | 0.07 | 62 | n.a. | n.a. | n.a. | 321 | 4.7 |
| cmpd 1.1 | 0.77 | 183 | 7.9 | 19 | AGO | 320 | 3.3 |
| cmpd 1.2 | 1.9 | 83 | n.a. | n.a. | n.a. | 355 | 2.2 |
| cmpd 1.3 | 1.8 | 254 | n.a. | n.a. | n.a. | 393 | 3.0 |
| cmpd 1.4 | 1.3 | 266 | n.a. | n.a. | n.a. | 324 | 1.6 |
| cmpd 2.1 | 1.2 | 143 | 0.56 | 52 | AGO | 304 | 3.2 |
| cmpd 2.2 | 10 | 110 | 4.0 | 103 | AGO | 344 | 3.7 |
| cmpd 2.3 | 6.1 | 88 | 1.2 | -7 | ANTAGO | 348 | 5.1 |
| cmpd 2.6 | 11 | 76 | 0.87 | 14 | AGO | 311 | 3.8 |
| cmpd 2.7 | 8.5 | 85 | 1.4 | 1 | ANTAGO | 318 | 5.0 |
| cmpd 2.8 | 9.4 | 126 | 2.0 | 87 | AGO | 298 | 3.2 |
| cmpd 2.9 | 4.8 | 70 | 2.4 | 19 | AGO | 347 | 3.2 |
| cmpd 2.10 | 5.8 | 128 | 2.8 | 30 | AGO | 324 | 2.9 |
| cmpd 2.11 | 18 | 85 | 3.6 | 57 | AGO | 316 | 2.5 |
| cmpd 2.12 | 12 | 109 | 4.5 | 45 | AGO | 351 | 4.0 |
| cmpd 3.2 | 13 | 116 | 2.2 | 32 | AGO | 359 | 4.0 |
| cmpd 3.3 | 12 | 173 | 4.0 | 102 | AGO | 286 | 4.1 |
| cmpd 3.4 | 7.9 | 158 | n.a. | n.a. | n.a. | 303 | 3.1 |
| cmpd 3.5 | 5.6 | 65 | n.a. | n.a. | n.a. | 326 | 3.9 |
| cmpd S01 | 6.0 | 37 | 0.4 | 103 | ANTAGO | 381 | 4.1 |
| cmpd S02 | 6.2 | 97 | 4.9 | 23 | AGO | 313 | 3.6 |
| cmpd S03 | 16 | 90 | 5.3 | 75 | AGO | 305 | 2.4 |
| cmpd S04 | 7.7 | 89 | 7.1 | 20 | AGO | 337 | 2.8 |

| | | | | | | | |
|---|---|---|---|---|---|---|---|
| cmpd S05 | 8.2 | 152 | 8.1 | 93 | ANTAGO | 330 | 3.7 |
| cmpd S06 | 2.9 | 76 | 10 | 25 | AGO | 306 | 2.7 |
| cmpd S07 | 20 | 149 | 12 | 21 | AGO | 308 | 3.3 |
| cmpd S08 | 11 | 181 | 24 | 37 | AGO | 301 | 2.9 |
| cmpd S09 | 3.1 | 144 | 38 | 37 | AGO | 346 | 2.9 |
| cmpd S10 | 9.6 | 116 | 43 | 92 | ANTAGO | 319 | 3.2 |
| cmpd S11 | 8.0 | 79 | 51 | 29 | AGO | 321 | 2.8 |
| cmpd S12 | 1.3 | 181 | n.a. | n.a. | n.a. | 327 | 2.5 |
| cmpd S13 | 1.5 | 128 | n.a. | n.a. | n.a. | 349 | 3.5 |
| cmpd S14 | 1.6 | 147 | n.a. | n.a. | n.a. | 288 | 2.7 |
| cmpd S15 | 1.6 | 157 | n.a. | n.a. | n.a. | 326 | 3.2 |
| cmpd S16 | 2.3 | 153 | n.a. | n.a. | n.a. | 298 | 3.0 |
| cmpd S17 | 2.5 | 69 | n.a. | n.a. | n.a. | 302 | 2.1 |
| cmpd S18 | 2.5 | 90 | n.a. | n.a. | n.a. | 323 | 2.9 |
| cmpd S19 | 2.8 | 170 | n.a. | n.a. | n.a. | 348 | 3.2 |
| cmpd S20 | 2.9 | 89 | n.a. | n.a. | n.a. | 356 | 3.0 |
| cmpd S21 | 3.5 | 169 | n.a. | n.a. | n.a. | 348 | 3.0 |
| cmpd S22 | 3.6 | 125 | n.a. | n.a. | n.a. | 349 | 1.6 |
| cmpd S23 | 3.6 | 162 | n.a. | n.a. | n.a. | 285 | 3.0 |
| cmpd S24 | 3.8 | 143 | n.a. | n.a. | n.a. | 352 | 4.2 |
| cmpd S25 | 4.3 | 82 | n.a. | n.a. | n.a. | 348 | 4.5 |
| cmpd S26 | 4.4 | 67 | n.a. | n.a. | n.a. | 397 | 4.6 |
| cmpd S27 | 4.6 | 152 | n.a. | n.a. | n.a. | 285 | 3.4 |
| cmpd S28 | 4.7 | 50 | n.a. | n.a. | n.a. | 352 | 3.2 |
| cmpd S29 | 4.8 | 94 | n.a. | n.a. | n.a. | 334 | 3.0 |
| cmpd S30 | 5.1 | 136 | n.a. | n.a. | n.a. | 319 | 3.0 |
| cmpd S31 | 5.1 | 81 | n.a. | n.a. | n.a. | 348 | 3.4 |

| | | | | | | | |
|---|---|---|---|---|---|---|---|
| cmpd S32 | 5.1 | 101 | n.a. | n.a. | n.a. | 351 | 4.4 |
| cmpd S33 | 5.1 | 166 | n.a. | n.a. | n.a. | 326 | 2.7 |
| cmpd S34 | 6.4 | 107 | n.a. | n.a. | n.a. | 282 | 2.0 |
| cmpd S35 | 6.6 | 82 | n.a. | n.a. | n.a. | 317 | 2.9 |
| cmpd S36 | 7.0 | 93 | n.a. | n.a. | n.a. | 373 | 3.8 |
| cmpd S37 | 7.1 | 163 | n.a. | n.a. | n.a. | 323 | 2.6 |
| cmpd S38 | 7.1 | 149 | n.a. | n.a. | n.a. | 321 | 2.8 |
| cmpd S39 | 7.6 | 161 | n.a. | n.a. | n.a. | 349 | 3.6 |
| cmpd S40 | 7.7 | 148 | n.a. | n.a. | n.a. | 313 | 2.6 |
| cmpd S41 | 7.8 | 152 | n.a. | n.a. | n.a. | 306 | 2.6 |
| cmpd S42 | 9.9 | 134 | n.a. | n.a. | n.a. | 369 | 3.0 |
| cmpd S43 | 11 | 84 | n.a. | n.a. | n.a. | 347 | 3.4 |
| cmpd S44 | 11 | 181 | n.a. | n.a. | n.a. | 321 | 3.3 |
| cmpd S45 | 11 | 89 | n.a. | n.a. | n.a. | 396 | 5.1 |
| cmpd S46 | 11 | 155 | n.a. | n.a. | n.a. | 332 | 4.1 |
| cmpd S47 | 11 | 123 | n.a. | n.a. | n.a. | 398 | 4.1 |
| cmpd S48 | 11 | 113 | n.a. | n.a. | n.a. | 320 | 4.2 |
| cmpd S49 | 12 | 160 | n.a. | n.a. | n.a. | 292 | 2.6 |
| cmpd S50 | 12 | 48 | n.a. | n.a. | n.a. | 329 | 5.1 |
| cmpd S51 | 12 | 171 | n.a. | n.a. | n.a. | 347 | 2.7 |
| cmpd S52 | 12 | 76 | n.a. | n.a. | n.a. | 312 | 1.0 |
| cmpd S53 | 12 | 163 | n.a. | n.a. | n.a. | 366 | 3.6 |
| cmpd S54 | 12 | 99 | n.a. | n.a. | n.a. | 367 | 2.5 |
| cmpd S55 | 13 | 180 | n.a. | n.a. | n.a. | 355 | 2.7 |
| cmpd S56 | 14 | 82 | n.a. | n.a. | n.a. | 377 | 4.4 |
| cmpd S57 | 14 | 113 | n.a. | n.a. | n.a. | 333 | 3.4 |
| cmpd S58 | 14 | 120 | n.a. | n.a. | n.a. | 329 | 2.4 |

| | | | | | | | |
|---|---|---|---|---|---|---|---|
| cmpd S59 | 15 | 100 | n.a. | n.a. | n.a. | 331 | 4.1 |
| cmpd S60 | 15 | 132 | n.a. | n.a. | n.a. | 356 | 3.7 |
| cmpd S61 | 16 | 71 | n.a. | n.a. | n.a. | 368 | 3.8 |
| cmpd S62 | 18 | 92 | n.a. | n.a. | n.a. | 339 | 3.2 |
| cmpd S63 | 18 | 158 | n.a. | n.a. | n.a. | 383 | 3.7 |
| cmpd S64 | 20 | 160 | n.a. | n.a. | n.a. | 333 | 2.3 |
| cmpd S65 | 21 | 125 | n.a. | n.a. | n.a. | 312 | 3.0 |
| cmpd S66 | 22 | 93 | n.a. | n.a. | n.a. | 295 | 2.5 |
| cmpd S67 | 23 | 126 | n.a. | n.a. | n.a. | 254 | 3.5 |
| cmpd S68 | 25 | 77 | n.a. | n.a. | n.a. | 381 | 3.4 |
| cmpd S69 | 27 | 156 | n.a. | n.a. | n.a. | 324 | 3.7 |
| cmpd S70 | 29 | 142 | n.a. | n.a. | n.a. | 323 | 1.0 |
| cmpd S71 | 30 | 160 | n.a. | n.a. | n.a. | 294 | 4.0 |
| cmpd S72 | 44 | 124 | n.a. | n.a. | n.a. | 371 | 3.6 |
| cmpd S73 | 44 | 128 | n.a. | n.a. | n.a. | 387 | 4.2 |
| cmpd S74 | 102 | 128 | n.a. | n.a. | n.a. | 389 | 4.2 |

[a] Efficacy relative to that of 100 nM fenoxycarb.

[b] Efficacy relative to 25 μM B[a]P in the agonist mode and 10 nM TCDD in the antagonist mode.

[c] AGO, agonist; ANTAGO, antagonist.

[d] Calculated lipophilicity.

n.a., not active.

#### References


1. Wen, Z., Zhang, Y., Zhang, B., Hang, Y., Xu, L., Chen, Y., Xie, Q., Zhao, Q., Zhang, L., Li, G. *et al.* (2023) Cryo-EM structure of the cytosolic AhR complex. *Structure* **31**, 295-308.e4.
2. Paramo, T., East, A., Garzón, D., Ulmschneider, M.B. and Bond, P.J. (2014) Efficient characterization of protein cavities within molecular simulation trajectories: trj_cavity. *J. Chem. Theory Comput*. **10**, 2151-2164.
3. Gruszczyk, J., Grandvuillemin, L., Lai-Kee-Him, J., Paloni, M., Savva, C.G., Germain, P., Grimaldi, M., Boulahtouf, A., Kwong, H.S., Bous, J. *et al.* (2022) Cryo-EM structure of the agonist-bound Hsp90-XAP2-AHR cytosolic complex. *Nat. Commun.* **13**, 7010.
